# Immunological memory to SARS-CoV-2 assessed for up to eight months after infection

**DOI:** 10.1101/2020.11.15.383323

**Authors:** Jennifer M. Dan, Jose Mateus, Yu Kato, Kathryn M. Hastie, Esther Dawen Yu, Caterina E. Faliti, Alba Grifoni, Sydney I. Ramirez, Sonya Haupt, April Frazier, Catherine Nakao, Vamseedhar Rayaprolu, Stephen A. Rawlings, Bjoern Peters, Florian Krammer, Viviana Simon, Erica Ollmann Saphire, Davey M. Smith, Daniela Weiskopf, Alessandro Sette, Shane Crotty

## Abstract

Understanding immune memory to SARS-CoV-2 is critical for improving diagnostics and vaccines, and for assessing the likely future course of the COVID-19 pandemic. We analyzed multiple compartments of circulating immune memory to SARS-CoV-2 in 254 samples from 188 COVID-19 cases, including 43 samples at ≥ 6 months post-infection. IgG to the Spike protein was relatively stable over 6+ months. Spike-specific memory B cells were more abundant at 6 months than at 1 month post symptom onset. SARS-CoV-2-specific CD4^+^ T cells and CD8^+^ T cells declined with a half-life of 3-5 months. By studying antibody, memory B cell, CD4^+^ T cell, and CD8^+^ T cell memory to SARS-CoV-2 in an integrated manner, we observed that each component of SARS-CoV-2 immune memory exhibited distinct kinetics.

---

Coronavirus disease 2019 (COVID-19), caused by the novel severe acute respiratory syndrome coronavirus 2 (SARS-CoV-2), is a serious disease that has resulted in widespread global morbidity and mortality. Humans make SARS-CoV-2-specific antibodies, CD4^+^ T cells, and CD8^+^ T cells in response to SARS-CoV-2 infection (*1–4*). Studies of acute and convalescent COVID-19 patients have observed that T cell responses are associated with reduced disease (*5–7*), suggesting that SARS-CoV-2-specific CD4^+^ T cell and CD8^+^ T cell responses may be important for control and resolution of primary SARS-CoV-2 infection. Ineffective innate immunity has been strongly associated with a lack of control of primary SARS-CoV-2 infection and a high risk of fatal COVID-19 (*8–12*), accompanied by innate cell immunopathology (*13–18*). Neutralizing antibodies have generally not correlated with lessened COVID-19 disease severity (*5*, *19, 20*), which was also observed for Middle Eastern respiratory syndrome (MERS), caused by MERS-CoV (*21*). Instead, neutralizing antibodies are associated with protective immunity against secondary infection with SARS-CoV-2 or SARS-CoV in non-human primates (*3*, *22*–*25*). Passive transfer of neutralizing antibodies in advance of infection (mimicking preexisting conditions upon secondary exposure) effectively limits upper respiratory tract (URT) infection, lower respiratory tract (lung) infection, and symptomatic disease in animal models (*26–28*). Passive transfer of neutralizing antibodies provided after initiation of infection in humans have had more limited effects on COVID-19 (*29*, *30*), consistent with a substantial role for T cells in control and clearance of an ongoing SARS-CoV-2 infection. Thus, studying antibody, memory B cell, CD4^+^ T cell, and CD8^+^ T cell memory to SARS-CoV-2 in an integrated manner is likely important for understanding the durability of protective immunity against COVID-19 generated by primary SARS-CoV-2 infection (*1*, *19*, *31*).

While sterilizing immunity against viruses can only be accomplished by high-titer neutralizing antibodies, successful protection against clinical disease or death can be accomplished by several other immune memory scenarios. Possible mechanisms of immunological protection can vary based on the relative kinetics of the immune memory responses and infection. For example, clinical hepatitis after hepatitis B virus (HBV) infection is prevented by vaccine-elicited immune memory even in the absence of circulating antibodies, because of the relatively slow course of HBV disease (*32*, *33*). The relatively slow course of severe COVID-19 in humans (median 19 days post-symptom onset (PSO) for fatal cases (*34*)) suggests that protective immunity against symptomatic or severe secondary COVID-19 may involve memory compartments such as circulating memory T cells and memory B cells (which can take several days to reactivate and generate recall T cell responses and/or anamnestic antibody responses) (*19*, *21*, *31*).

Immune memory, from either primary infection or immunization, is the source of protective immunity from a subsequent infection (*35*–*37*). Thus, COVID-19 vaccine development relies on immunological memory (*1*, *3*). Despite intensive study, the kinetics, duration, and evolution of immune memory in humans to infection or immunization are not in general predictable based on the initial effector phase, and immune responses at short time points after resolution of infection are not very predictive of long-term memory (*38*–*40*). Thus, assessing responses over an interval of six months or more is usually required to ascertain the durability of immune memory.

A thorough understanding of immune memory to SARS-CoV-2 requires evaluation of its various components, including B cells, CD8^+^ T cells, and CD4^+^ T cells, as these different cell types may have immune memory kinetics relatively independent of each other. Understanding the complexities of immune memory to SARS-CoV-2 is key to gain insights into the likelihood of durability of protective immunity against re-infection with SARS-CoV-2 and secondary COVID-19 disease. In the current study, we assessed immune memory of all three branches of adaptive immunity (CD4^+^ T cell, CD8^+^ T cell, and humoral immunity) in a predominantly cross-sectional study of 188 recovered COVID-19 cases, extending up to eight months post-infection. The findings have implications for immunity against secondary COVID-19, and thus the potential future course of the pandemic (*41, 42).*

## COVID-19 cohort

188 individuals with COVID-19 were recruited for this study. Subjects (80 male, 108 female) represented a range of asymptomatic, mild, moderate, and severe COVID-19 cases (**Table 1**), and were recruited from multiple sites throughout the United States. The majority of subjects were from California or New York. Most subjects had a “mild” case of COVID-19, not requiring hospitalization. 93% of subjects were never hospitalized for COVID-19; 7% of subjects were hospitalized, some of whom required intensive care unit (ICU) care (**Table 1**). This case severity distribution was consistent with the general distribution of symptomatic disease severity among COVID-19 cases in the USA. The study primarily consisted of symptomatic disease cases (97%, **Table 1**), due to the nature of the study recruitment design. Subject ages ranged from 19 to 81 years old (**Table 1**). Most subjects provided a blood sample at a single time point, between 6 days post-symptom onset (PSO) and 240 days PSO (**Table 1**), with 43 samples at ≥ 6 months PSO (178 days or longer). Additionally, 51 subjects in the study provided longitudinal blood samples over a duration of several months (2-4 time points; **Table 1**), allowing for longitudinal assessment of immune memory in a subset of the cohort.

**Table 1.**
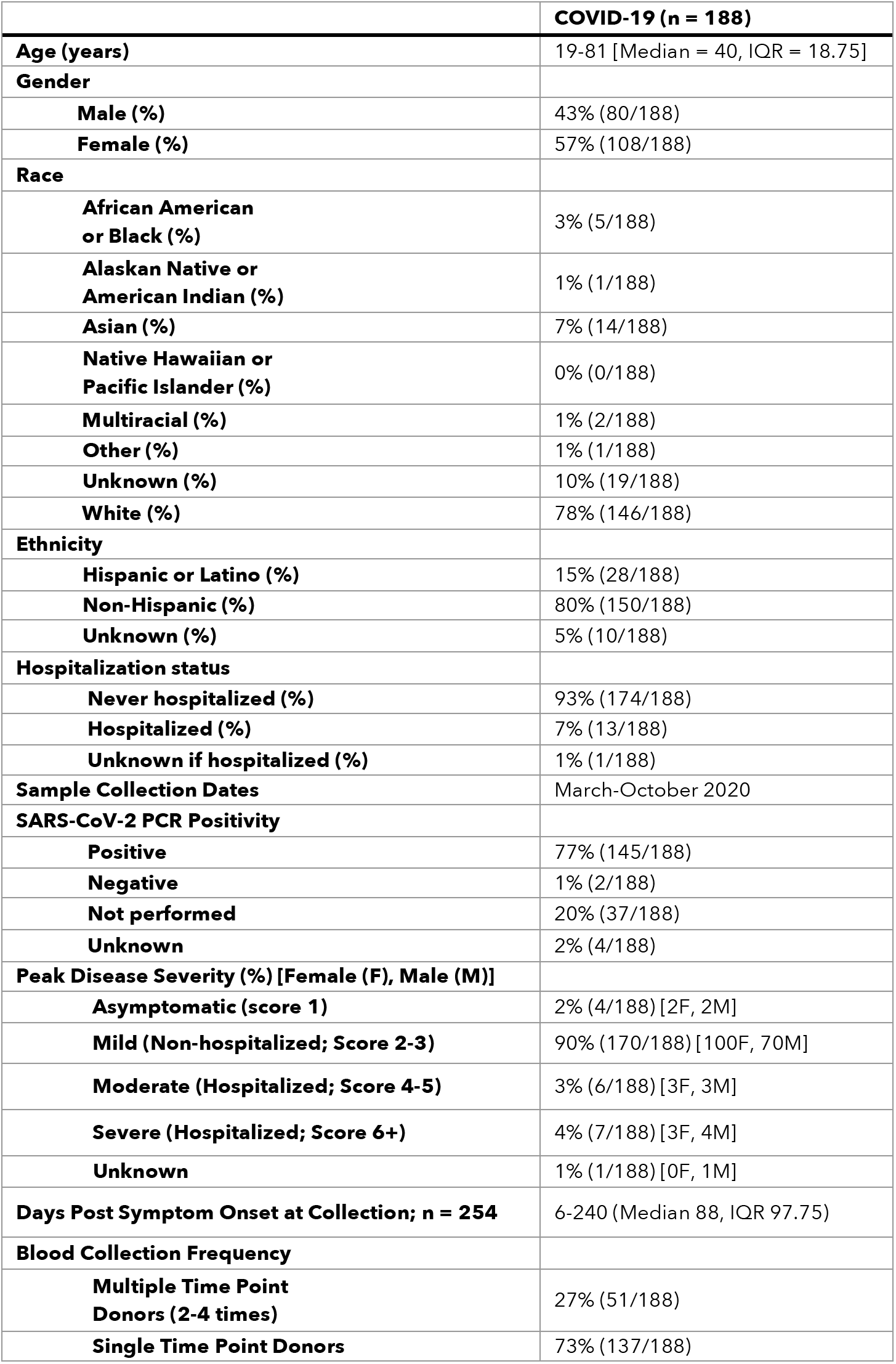
Participant characteristics

## SARS-CoV-2 circulating antibodies over time

The vast majority of SARS-CoV-2 infected individuals seroconvert, at least for a duration of months (*1*, *2*, *4*, *43–45*). Seroconversion rates range from 91-99% in large studies (*44*, *45*). Durability assessments of circulating antibody titers in Figure 1 were based on data ≥ 20 days PSO, with the plot of the best fitting curve fit model shown in blue (see Methods). SARS-CoV-2 Spike immunoglobulin G (IgG) endpoint ELISA titers in plasma were measured for all subjects of this cohort (**Fig. 1A-B**). Spike receptor binding domain (RBD) IgG was also measured (**Fig. 1C-D**), as RBD is the target of most neutralizing antibodies against SARS-CoV-2 (*4*, *27*, *46, 47*). SARS-CoV-2 pseudovirus (PSV) neutralizing antibody titers were measured in all subjects (**Fig. 1E-F**). Nucleocapsid (N) IgG endpoint ELISA titers were also measured for all subjects (**Fig. 1G-H**), as Nucleocapsid is a common antigen in commercial SARS-CoV-2 serological test kits.

**Fig. 1.**
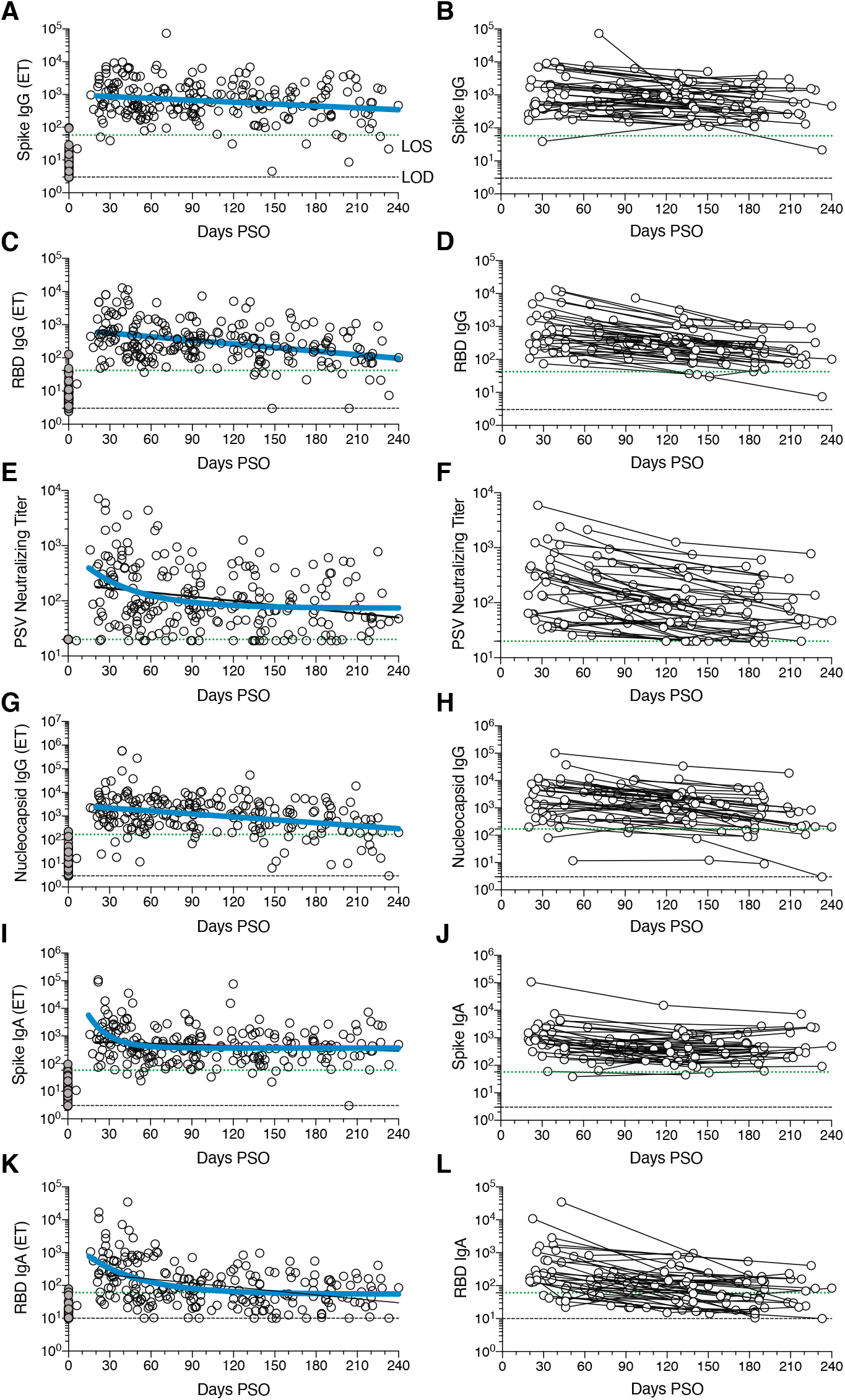
Circulating antibodies to **SARS-CoV-2 over time.** **(A)** Cross-sectional Spike IgG from COVID-19 subject plasma samples (n=228). Continuous decay preferred model for best fit curve, *t*_1/2_ = 140 days, 95% CI: 89-325 days. R = −0.23, p=0.0006. **(B)** Longitudinal Spike IgG (n=51), average *t*_1/2_ = 103 days, 95% CI: 65-235 days **(C)** Cross-sectional RBD IgG. Continuous decay preferred model for best fit curve, *t*_1/2_ = 83 days, 95% CI: 62 to 126 days. R = −0.36, p<0.0001. **(D)** Longitudinal RBD IgG, average *t*_1/2_ = 69 days, 95% CI: 58-87 days **(E)** Cross-sectional SARS-CoV-2 PSV neutralizing titers. One-phase decay (blue line) preferred model for best fit curve, initial *t*_1/2_ = 27 days, 95% CI 11-157d. R = −0.32. Continuous decay fit line shown as black line. **(F)** Longitudinal PSV neutralizing titers of SARS-CoV-2 infected subjects, average *t*_1/2_ = 90 days, 95% CI: 70-125 days. **(G)** Cross-sectional Nucleocapsid IgG. Continuous decay preferred model for best fit curve, *t*_1/2_ = 68 days, 95% CI: 50-106 days. R = −0.34, p<0.0001. **(H)** Longitudinal Nucleocapsid IgG, average *t*_1/2_ = 68 days, 95% CI: 55-90 days. **(I)** Crosssectional Spike IgA titers. One-phase decay (blue line) preferred model for best fit curve, initial *t*_1/2_ = 11 days, 95% CI 5-25d. R = −0.30. Continuous decay fit shown as black line. **(J)** Longitudinal Spike IgA, *t*_1/2_ = 210 days, 95% CI 126-627 days. **(K)** Cross-sectional RBD IgA. One-phase decay (blue line) preferred model for best fit curve, initial *t*_1/2_ = 27 days, 95% CI: 15-59 days. R = −0.45. Continuous decay line fit shown in black. **(L)** Longitudinal RBD IgA, average *t*_1/2_ = 74 days, 95% CI: 56-107 days. For cross-sectional analyses, SARS-CoV-2 infected subjects (white circles, n=238) and unexposed subjects (gray circles, n=51). For longitudinal samples, SARS-CoV-2 subjects (n=51). The dotted black line indicates limit of detection (LOD). The dotted green line indicates limit of sensitivity (LOS) above uninfected controls. Unexposed = gray, COVID subjects = white. Log data analyzed in all cases. Thick blue line represents best fit curve. When two fit curves are shown, the thin black line represents the alternative fit curve.

SARS-CoV-2 Spike IgG titers were relatively stable from 20-240 days PSO, when assessing all COVID-19 subjects by cross-sectional analysis (half-life *t*_1/2_ = 140 days, **Fig. 1A**). Spike IgG titers were heterogenous among subjects (range 5 to 73,071; 575 median), as has been widely observed (*45*, *47*). This gave a wide confidence interval for the Spike IgG *t*_1/2_ (95% CI: 89 to 325 days). While the antibody responses may have more complex underlying decay kinetics, the best fit curve was a continuous decay, likely related to heterogeneity between individuals. SARS-CoV-2 Nucleocapsid IgG kinetics were similar to Spike IgG over 8 months (*t*_1/2_ 68 days, 95% CI: 50-106 days. **Fig. 1G**). As a complementary approach, using paired samples from the subset of subjects who donated at two or more time points, the calculated Spike IgG titer average *t*_1/2_ was 103 days, (95% CI: 66-235 days, **Fig. 1B**) and the Nucleocapsid IgG titer average *t*_1/2_ was 68 days, (95% CI: 55-90 days, **Fig. 1H**). The percentage of subjects seropositive for Spike IgG at 1 month PSO (20-50 days) was 98% (54/55). The percentage of subjects seropositive for Spike IgG at 6 to 8 months PSO (≥178 days) was 90% (36/40).

Cross-sectional analysis of SARS-CoV-2 RBD IgG titers from 20-240 days PSO gave an estimated *t*_1/2_ of 83 days (95% CI: 62-126 days, **Fig. 1C**). As a complementary approach, we again used paired samples, which gave an average *t*_1/2_ of 69 days (95% CI: 58-87 days, **Fig. 1D**). The percentage of subjects seropositive for RBD IgG at 6 to 8 months PSO was 88% (35/40). Thus, RBD IgG titer maintenance largely matched that of Spike IgG. SARS-CoV-2 PSV neutralization titers in the full cohort largely matched the results of SARS-CoV-2 RBD IgG ELISA binding titers (**Fig.1 E-F**). A one-phase decay model was the best fit (P=0.015, F test. Initial decay *t*_1/2_ 27 days, followed by an extended plateau phase. **Fig. 1E**), while a continuous decay fit gave an estimated *t*_1/2_ of 114 days (**Fig. 1E,** black line). Paired timepoints analysis of the PSV neutralization titers gave an estimated *t*_1/2_ of 90 days, (95% CI: 70-125 days, **Fig. 1F**). The percentage of subjects seropositive for SARS-CoV-2 neutralizing antibodies (titer ≥ 20) at 6 to 8 months PSO was 90% (36/40). Notably, even low levels of circulating neutralizing antibody titers (≥ 1:20) were associated with a substantial degree of protection against COVID-19 in non-human primates (*24*, *48*). Thus, modest levels of circulating SARS-CoV-2 neutralizing antibodies are of biological interest in humans.

SARS-CoV-2 Spike IgA (**Fig. 1I-J**) and RBD IgA (**Fig.1K-L**) titers were also assessed. Paired timepoints analysis of Spike IgA titers yielded an estimated *t*_1/2_ of 210 days (95% CI 126-703 days, **Fig. 1J**). Cross-sectional analysis of Spike IgA fit a short one-phase decay model with an extended plateau phase (initial *t*_1/2_ of 14 days, **Fig. 1I**). Circulating RBD IgA had an estimated initial *t*_1/2_ of 27 days, decaying by ~90 days in most COVID-19 cases to levels indistinguishable from uninfected controls (**Fig. 1K**), consistent with observations 3 months PSO (*44*, *49*). By paired sample analysis, long-lasting RBD IgA was made in some subjects, but often near the limit of sensitivity (LOS) (**Fig. 1L**).

## SARS-CoV-2 memory B cells

To identify SARS-CoV-2-specific memory B cells, fluorescently labeled multimerized probes were used to detect B cells specific to Spike, RBD, and Nucleocapsid (**Fig 2A,** Fig. S1). Antigen-binding memory B cells (defined as IgD^-^ and/or CD27^+^) were further distinguished according to surface Ig isotypes: IgM, IgG or IgA (**Fig. 2B,** Fig. S1).

**Fig. 2.**
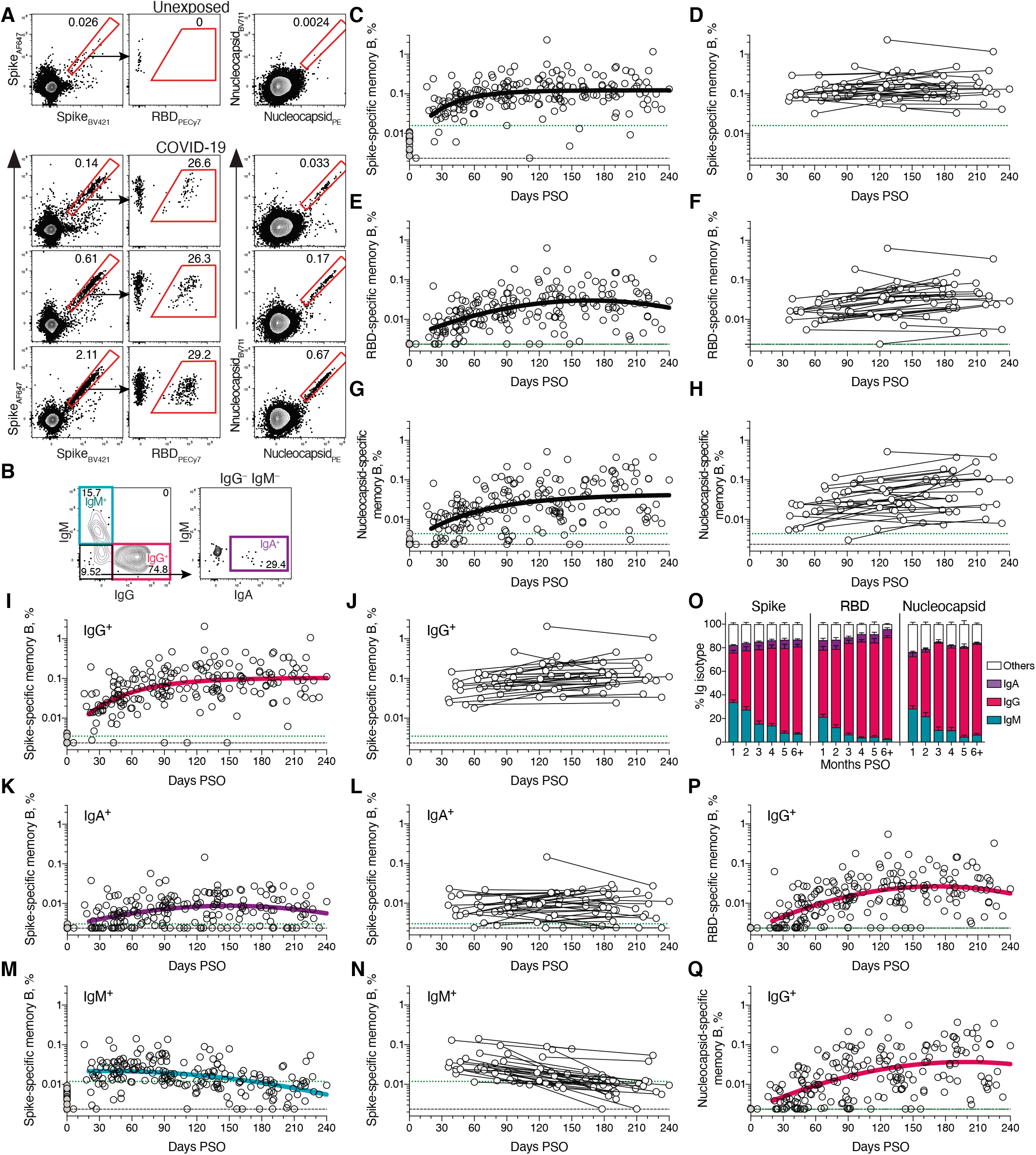
Kinetics of SARS-CoV-2 memory B cell responses. (**A**) Example flow cytometry plots showing staining patterns of SARS-CoV-2 antigen probes on memory B cells (See Fig. S1 for gating). One unexposed donor and three convalescent COVID-19 subjects are shown. Numbers indicate percentages. (**B**) Gating strategies to define IgM^+^, IgG^+^, or IgA^+^ SARS-CoV-2 Spike-specific memory B cells. The same gating strategies were used for RBD- or Nucleocapsid-specific B cells. (**C**) Crosssectional analysis of frequency (% of CD19^+^ CD20^+^ B cells) of SARS-CoV-2 S-specific total (IgG^+^, IgM^+^, or IgA^+^) memory B cells. Pseudo-first order kinetic model for best fit curve (R = 0.38). (**D**) Longitudinal analysis of SARS-CoV-2 Spike-specific memory B cells. (**E**) Cross-sectional analysis of SARS-CoV-2 RBD-specific total (IgG^+^, IgM^+^, or IgA^+^) memory B cells. Second order polynomial model for best fit curve (R = 0.46). (**F**) Longitudinal analysis of SARS-CoV-2 RBD-specific memory B cells. (**G**) Cross-sectional analysis of SARS-CoV-2 Nucleocapsid-specific total (IgG^+^, IgM^+^, or IgA^+^) memory B cells. Pseudo-first order kinetic model for best fit curve (R = 0.44). (**H**) Longitudinal analysis of IgG^+^ SARS-CoV-2 Nucleocapsid-specific memory B cells. (**I**) Cross-sectional analysis of SARS-CoV-2 Spike-specific IgG^+^ memory B cells. Pseudo-first order kinetic model for best fit curve (R = 0.49). (**J**) Longitudinal analysis of SARS-CoV-2 Spike-specific IgG^+^ memory B cells. (**K**) Cross-sectional analysis of SARS-CoV-2 Spikespecific IgA^+^ memory B cells. Second order polynomial model for best fit curve (|R| = 0.32). (**L**) Longitudinal analysis of SARS-CoV-2 Spike-specific IgA^+^ memory B cells. (**M**) Cross-sectional analysis of SARS-CoV-2 Spike-specific IgM^+^ memory B cells. Second order polynomial model for best fit curve (|R| = 0.41). (**N**) Longitudinal analysis of SARS-CoV-2 Spike-specific IgM^+^ memory B cells. (**O**) Fraction of SARS-CoV-2 antigen-specific memory B cells that belong to indicated Ig isotypes at 1-8 months PSO. Mean ± SEM. (**P**) Cross-sectional analysis of SARS-CoV-2 RBD-specific IgG^+^ memory B cells. Second order polynomial model for best fit curve (|R| = 0.51). (**Q**) Cross-sectional analysis of SARS-CoV-2 Nucleocapsid-specific IgG^+^ memory B cells. Second order polynomial model for best fit curve (|R| = 0.51). n = 20 unexposed subjects (gray circles) and n = 160 COVID-19 subjects (n = 197 data points, white circles) for cross-sectional analysis. n = 36 COVID-19 subjects (n = 73 data points, white circles) for longitudinal analysis. The dotted black line indicates limit of detection (LOD). The dotted green line indicates limit of sensitivity (LOS).

Cross-sectional analysis of COVID-19 subjects revealed that frequencies of SARS-CoV-2 Spike-specific memory B cells increased over the first ~120 days PSO and then plateaued (pseudo-first order model for best fit curve, R = 0.38. Better fit than second order polynomial model by Akaike’s Information Criterion. **Fig 2C,** Fig. S2A). Spike-specific memory B cell frequencies increased from the first time-point (36-163 days) to the second time-point (111-240 days) in paired samples from 24 of 36 longitudinally tracked donors (**Fig 2D**). Spike-specific memory B cells in SARS-CoV-2-unexposed subjects were rare (median 0.0078%. **Fig 2A, 2C**).

RBD-specific memory B cells displayed similar kinetics to Spike-specific memory B cells. RBD-specific memory B cells were undetectable in SARS-CoV-2 unexposed subjects (**Fig. 2E.** Fig. S2C), as expected. RBD-specific memory B cells appeared as early as 16 days PSO, and the frequency steadily increased in the following 4-5 months (**Fig. 2E.** Fig. S2B-C). 29 of 36 longitudinally tracked individuals had higher frequencies of RBD-specific memory B cells at the later time point (**Fig. 2F**), again showing an increase in SARS-CoV-2 specific memory B cells several months post-infection. ~10-30% of Spike-specific memory B cells from SARS-CoV-2 convalescent donors were specific for the RBD domain (**Fig. 2A,** Fig. S2B).

SARS-CoV-2 Nucleocapsid-specific memory B cells were also detected after SARS-CoV-2 infection (**Fig. 2A**). Similar to Spike- and RBD-specific memory B cells, Nucleocapsid-specific memory B cell frequency steadily increased during the first ~4-5 months PSO (**Fig. 2G, 2H,** Fig. S2D). Antibody affinity maturation could potentially explain the increased frequencies of SARS-CoV-2-specific memory B cells detected by the antigen probes. However, geometric mean fluorescent intensity (MFI) of probe binding was stable over time (Fig. S2I-J), not supporting an affinity maturation explanation for the increased memory B cell frequencies.

Representation of Ig isotypes among the SARS-CoV-2 Spike-specific memory B cell population shifted with time (**Fig. 2I-2O**). During the earliest phase of memory (20-60 days PSO), IgM^+^ and IgG^+^ isotypes were similarly represented (**Fig. 2O**), but IgM^+^ memory B cells then declined (**Fig. 2M-O**), and IgG^+^ Spike-specific memory B cells then dominated by 6 months PSO (**Fig. 2O**). IgA^+^ Spike-specific memory B cells were detected as a small fraction of the total Spike-specific memory B cells (~5%, **Fig. 2O**). IgG^+^ Spike-specific memory B cell frequency increased while IgA^+^ was low and stable over the 8 months period (**Fig. 2I-2L**). Similar patterns of increasing IgG^+^ memory, short-lived IgM^+^ memory, and stable IgA^+^ memory were observed for RBD- and Nucleocapsid-specific memory B cells over the 8 months period (**Fig. 2O-2Q, Fig. S2E-S2H**).

There is limited knowledge of memory B cell kinetics following primary acute viral infection in humans. A recently published SARS-CoV-2 study found RBD-specific memory B cells out to ~90 days PSO, with increasing frequencies (and a low frequency of IgA^+^ cells) (*50*), consistent with observations reported here. For other acute infectious diseases, we are not currently aware of other cross-sectional or longitudinal analyses of antigen-specific memory B cells by flow cytometry covering a 6+ month window after infection, except for four individuals with Ebola (*51*) and two individuals studied after yellow fever virus immunization (*52*) (we exclude influenza vaccines for comparison here, because people have numerous exposures and complex immune history to influenza). In the yellow fever study, short-lived IgM^+^ memory and longer-lasting isotype-switched memory B cells were observed in the two individuals. Overall, based on the observations here, development of B cell memory to SARS-CoV-2 was robust, and is likely long-lasting.

## SARS-CoV-2 memory CD8^+^ T cells

SARS-CoV-2 memory CD8^+^ T cells were measured in 169 COVID-19 subjects using a series of 23 peptide pools covering the entirety of the SARS-CoV-2 ORFeome (*2*, *5*). The most commonly recognized ORFs were Spike, Membrane (M), Nucleocapsid, and ORF3a (CD69^+^ CD137^+^, **Fig. 3A** and Fig. S3A-B), consistent with our previous study (*2*). The percentage of subjects with detectable circulating SARS-CoV-2 memory CD8^+^ T cells at 1 month PSO (20-50 days) was 70% (40/57, **Fig. 3B**). The proportion of subjects positive for SARS-CoV-2 memory CD8^+^ T cells at ≥ 6 months PSO was 50% (18/36). This could potentially underestimate CD8^+^ T cell memory, as 15-mers can be suboptimal for detection of some antigen-specific CD8^+^ T cells (*53*); however, pools of predicted SARS-CoV-2 class I epitope of optimal size also detected virus-specific CD8^+^ T cells in ~70% of individuals 1-2 months PSO, indicating consistency between the two experimental approaches (*2*).

**Fig. 3.**
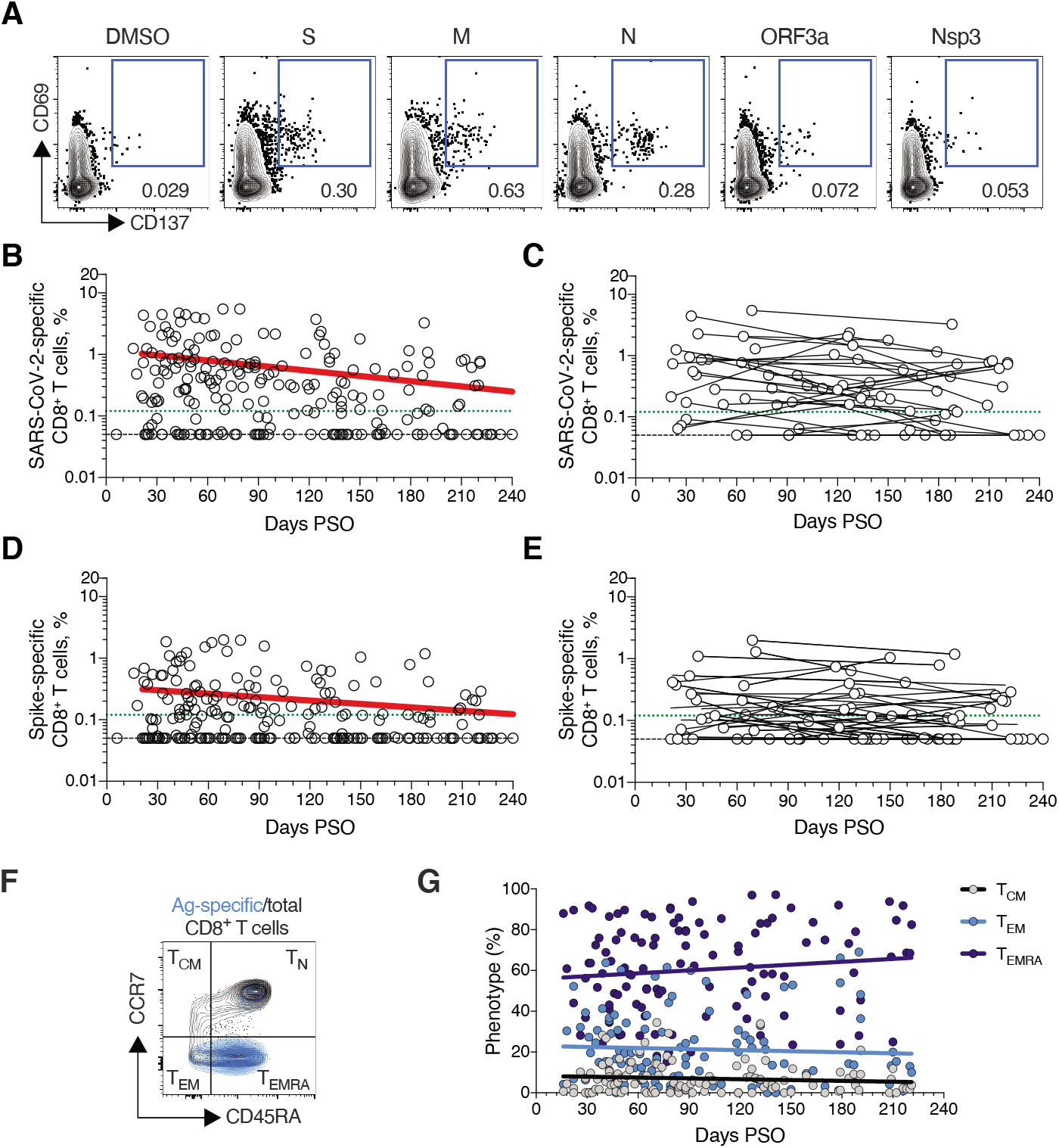
SARS-CoV-2 circulating memory CD8^+^ T cells. **(A)** Representative flow cytometry plots of SARS-CoV-2-specific CD8^+^ T cells (CD69^+^ CD137^+^, See Fig. S3 for gating) after overnight stimulation with S, N, M, ORF3a, or nsp3 peptide pools, compared to negative control (DMSO). **(B)** Cross-sectional analysis of frequency (% of CD8^+^ T cells) of total SARS-CoV-2-specific CD8^+^ T cells. Continuous decay preferred fit model, t1/2 = 125 days. R = −0.24, p = 0.0003. **(C)** Longitudinal analysis of total SARS-CoV-2-specific CD8^+^ T cells in paired samples. **(D)** Cross-sectional analysis of Spike-specific CD8^+^ T cells. Linear decay preferred model, t1/2 = 225 days. R = −0.18, p = 0.007. **(E)** Longitudinal analysis of Spikespecific CD8^+^ T cells in paired samples. **(F)** Distribution of central memory (T_CM_), effector memory (T_EM_), and terminally differentiated effector memory cells (T_EMRA_) among total SARS-CoV-2-specific CD8^+^ T cells. n = 169 COVID-19 subjects (n = 215 data points, white circles) for cross-sectional analysis. n = 37 COVID-19 subjects (n = 83 data points, white circles) for longitudinal analysis. The dotted black line indicates limit of detection (LOD). The dotted green line indicates limit of sensitivity (LOS).

SARS-CoV-2 memory CD8^+^ T cells declined with an apparent *t*_1/2_ of 125 days in the full cohort (**Fig. 3B**) and *t*_1/2_ 190 days among 29 paired samples (**Fig. 3C**). Spike-specific memory CD8^+^ T cells exhibited similar kinetics to the overall SARS-CoV-2-specific memory CD8^+^ T cells (*t*_1/2_ 225 days for the full cohort and 185 days among paired samples, **Fig. 3D-E**, respectively). Phenotypic markers indicated that the majority of SARS-CoV-2-specific memory CD8^+^ T cells were terminally differentiated effector memory cells (T_EMRA_) (*54*), with small populations of central memory (T_CM_) and effector memory (TEM) (**Fig. 3F**). In the context of influenza, CD8^+^ T_EMRA_ cells were associated with protection against severe disease in humans (*55*). The memory CD8^+^ T cell half-lives observed here were comparable to the 123 days *t*_1/2_ observed for memory CD8^+^ T cells after yellow fever immunization (*56*). Thus, the kinetics of circulating SARS-CoV-2-specific CD8^+^ T cell were consistent with what has been reported for another virus that causes acute infections in humans.

## SARS-CoV-2 memory CD4^+^ T cells

SARS-CoV-2 memory CD4^+^ T cells were identified in 169 subjects using the same series of 23 peptide pools covering the SARS-CoV-2 ORFeome (*2*, *5*). The most commonly recognized ORFs were Spike, M, Nucleocapsid, ORF3a, and nsp3 (CD137^+^ OX40^+^, **Fig. 4A** and Fig. S4A-B), consistent with our previous study (*2*). Circulating SARS-CoV-2 memory CD4^+^ T cell responses were quite robust (**Fig. 4B**); 42% (24/57) of COVID-19 cases at 1 month PSO had > 1.0% SARS-CoV-2-specific CD4^+^ T cells. SARS-CoV-2 memory CD4^+^ T cells declined with an apparent *t*_1/2_ of 94 days in the full cohort (**Fig. 4B**) and *t*_1/2_ 64 days among 36 paired samples (**Fig. 4C**). The percentage of subjects with detectable circulating SARS-CoV-2 memory CD4^+^ T cells at 1 month PSO (20-50 days) was 93% (53/57, **Fig. 4B**). The proportion of subjects positive for SARS-CoV-2 memory CD4^+^ T cells at ≥ 6 months PSO was 92% (33/36).

**Fig. 4.**
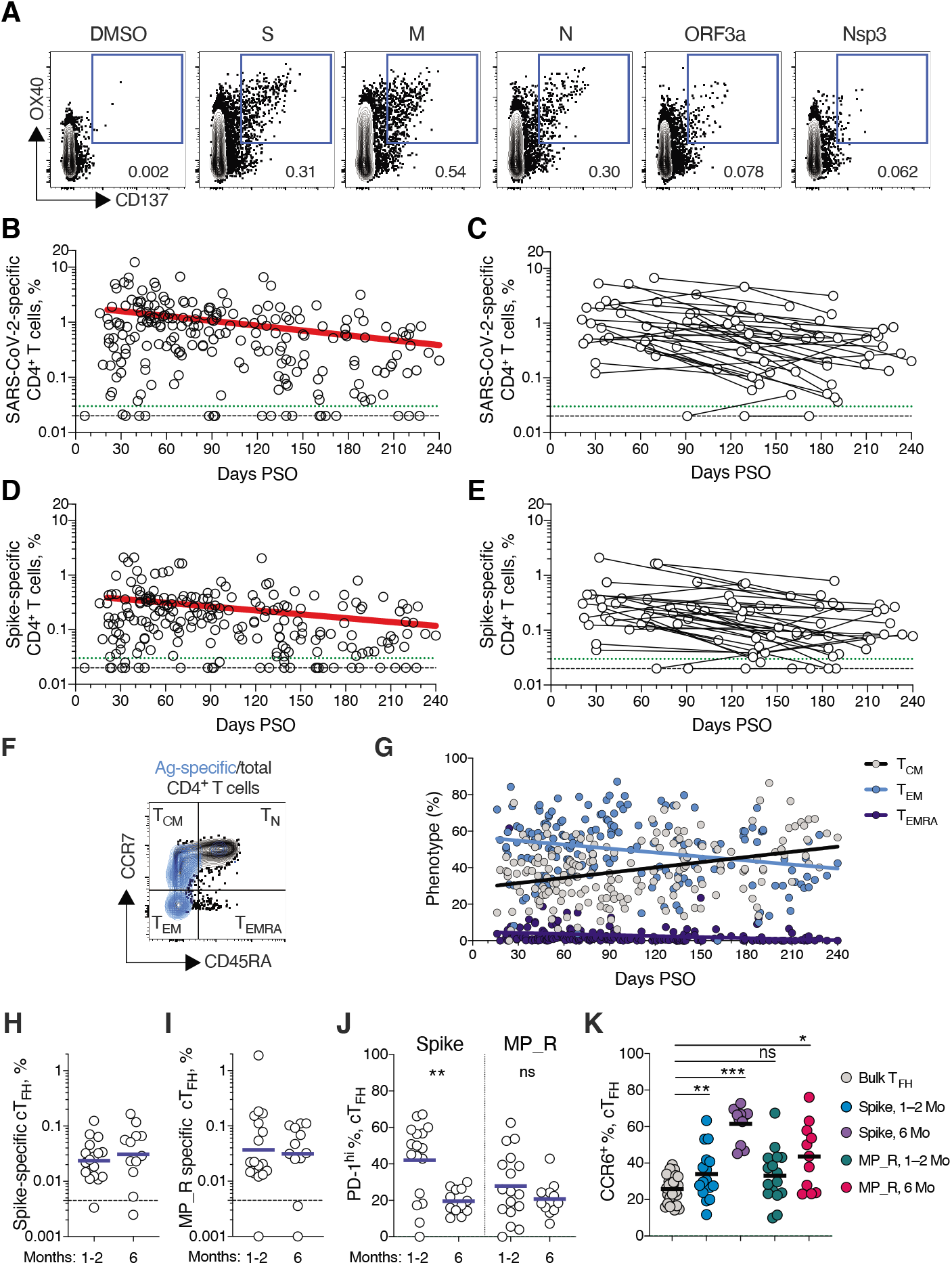
SARS-CoV-2 circulating memory CD4^+^ T cells. **(A)** Representative flow cytometry plots of SARS-CoV-2-specific CD4^+^ T cells (CD137^+^ OX40^+^, See Fig S4 for gating) after overnight stimulation with S, N, M, ORF3a, or nsp3 peptide pools, compared to negative control (DMSO). **(B)** Cross-sectional analysis of frequency (% of CD4^+^ T cells) of total SARS-CoV-2-specific CD4^+^ T cells. Continuous decay preferred fit model, *t*_1/2_ = 94 days. R = −0.29, p<0.0001. **(C)** Longitudinal analysis of total SARS-CoV-2-specific CD4^+^ T cells in paired samples from the same subjects. **(D)** Cross-sectional analysis of Spikespecific CD4^+^ T cells. Linear decay preferred model, t1/2 = 139 days. R = −0.26, p<0.0001. **(E)** Longitudinal analysis of Spike-specific CD4^+^ T cells in paired samples from the same subjects. **(F, G)** Distribution of central memory (T_CM_), effector memory (T_EM_), and terminally differentiated effector memory cells (T_EMRA_) among total SARS-CoV-2-specific CD4^+^ T cells. (**H, I**) Quantitation of SARS-CoV-2-specific circulating T follicular helper (cT_FH_) cells (surface CD40L^+^ OX40^+^, as % of CD4^+^ T cells. See Fig S5 for gating) after overnight stimulation with **(H)** Spike (S) or **(I)** MP_R peptide pools. (**J**) PD-1^hi^ SARS-CoV-2-specific T_FH_ at 1-2 months (mo) and 6 mo PSO. (**K**) CCR6^+^ SARS-CoV-2-specific cT_FH_ in comparison to bulk cT_FH_ cells in blood. For (**A-E**), n = 169 COVID-19 subjects (n = 215 data points, white circles) for cross-sectional analysis, n = 37 COVID-19 subjects (n = 83 data points, white circles) for longitudinal analysis. The dotted black line indicates limit of detection (LOD). The dotted green line indicates limit of sensitivity (LOS). For **(H-J**), n = 29 COVID-19 subject samples (white circles), n = 17 COVID-19 subjects at 1-2 mo, n = 12 COVID-19 subjects at 6 mo. The dotted black line indicates limit of detection (LOD). Statistics by (**J**) Mann-Whitney U test and (**K**) Wilcoxon signed-rank test. * p<0.05, **p<0.01, *** p<0.001.

Spike-specific and M-specific memory CD4^+^ T cells exhibited similar kinetics to the overall SARS-CoV-2-specific memory CD4^+^ T cells (whole cohort *t*_1/2_ 139 days and 153 days, respectively. **Fig. 4D-E**, and Fig. S4D). A plurality of the SARS-CoV-2 memory CD4^+^ T cells present at ≥ 6 months PSO had a T_CM_ phenotype (**Fig. 4F**).

T follicular helpers (T_FH_) are the specialized subset of CD4^+^ T cells required for B cell help (*57*), and are therefore critical for the generation of neutralizing antibodies and long-lived humoral immunity in most contexts. Thus, we examined circulating T_FH_ (cT_FH_) memory CD4^+^ T cells, with particular interest in Spike-specific memory cT_FH_ cells due to the importance of antibody responses against Spike. Memory cT_FH_ cells specific for predicted epitopes across the remainder of the SARS-CoV-2 genome were also measured, using the MP_R megapool. Memory cT_FH_ cells specific for SARS-CoV-2 Spike and MP_R were detected in the majority of COVID-19 cases at early time points (16/17. **Fig. 4H-I**, and Fig. S5A-D). cT_FH_ memory appeared to be stable, with almost all subjects positive for Spike and MP_R memory cT_FH_ cells at 6 months PSO (11/12 & 10/12, respectively. **Fig. 4H-I**). Recently activated cT_FH_ cells are PD-1^hi^ (*57*). Consistent with conversion to resting memory cT_FH_ cells, the percentage of PD-1^hi^ SARS-CoV-2-specific memory cT_FH_ dropped over time (**Fig. 4J**). CCR6^+^ SARS-CoV-2-specific cT_FH_ cells have been associated with reduced COVID-19 disease severity (*5*) and have been reported to be a major fraction of Spike-specific cT_FH_ cells in some studies (*5*, *50*, *58*). Here we confirmed that a significant fraction of both Spike-specific and MP_R memory cT_FH_ cells were CCR6^+^. We also observed increases in CCR6^+^ cT_FH_ memory over time (p=0.001 and p=0.014 at ≥ 6 months PSO compared to bulk cT_FH_. **Fig. 4K**). Overall, substantial cT_FH_ memory was observed after SARS-CoV-2 infection, with durability ≥ 6 months PSO.

## Immune memory relationships

Immune memory to SARS-CoV-2 were considered, including relationships between the compartments of immune memory. Males had higher Spike IgG (ANCOVA p=0.00018, **Fig. 5A**) and RBD and Nucleocapsid IgG (ANCOVA p=0.00077 & p=0.018, Fig. S6A-B), consistent with other studies (*46*, *47*). Higher Spike IgG was also observed in males when only non-hospitalized cases were considered (ANCOVA p=0.00025, Fig. S6C). In contrast, no differences were observed in IgA or PSV neutralization titers (Fig. S6D-F), and no differences were detected in SARS-CoV-2 memory B cell, memory CD8^+^ T cell, or memory CD4^+^ T cell frequencies between males and females (Fig. S6G-K).

**Fig. 5.**
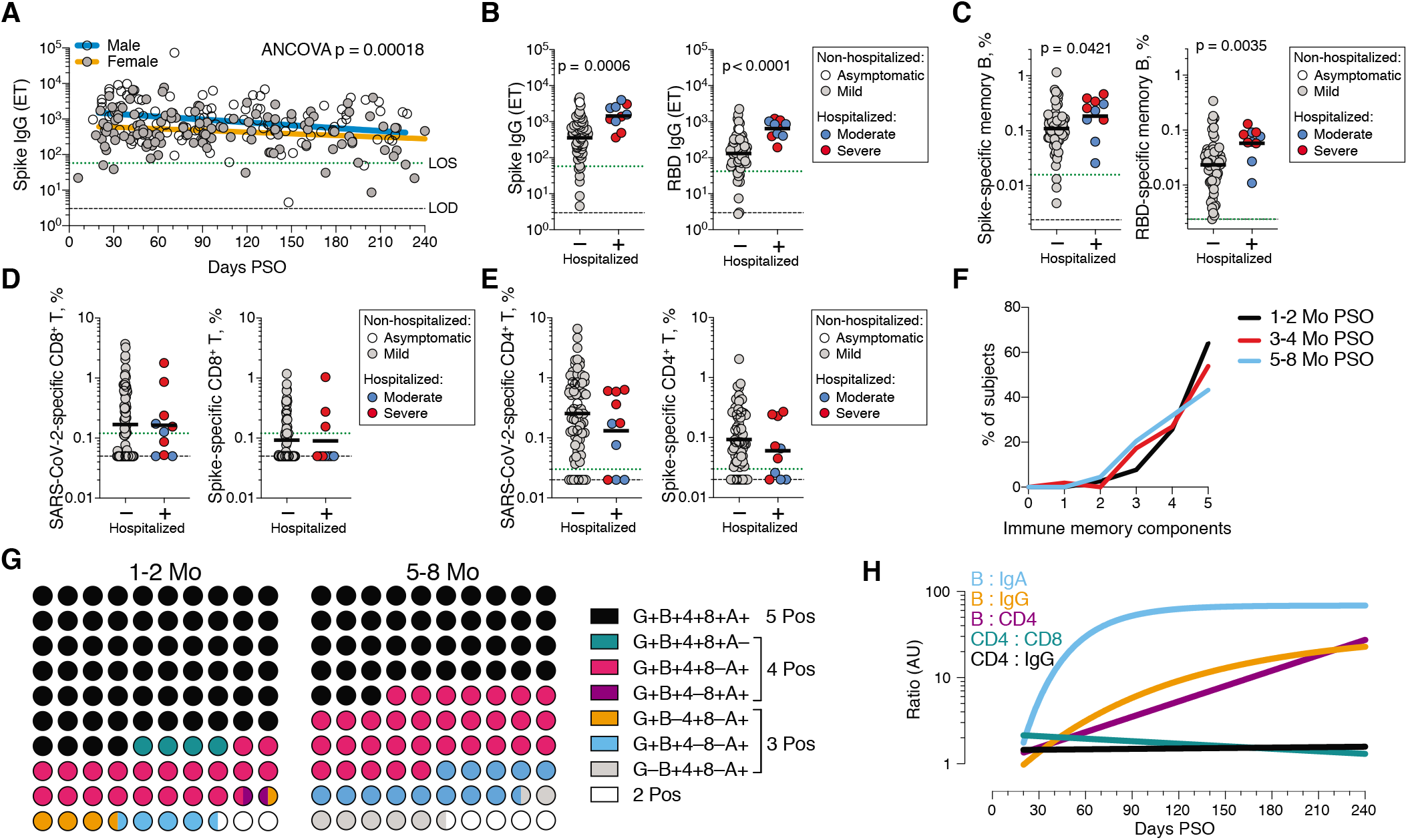
Immune memory relationships. **(A)** Relationship between gender and Spike IgG titers over time. Males: Linear decay preferred model, *t*_1/2_ = 110 days, 95% CI: 65-349 days, R = −0.27, p = 0.0046. Females: linear decay preferred model, *t*_1/2_ = 159 days, 95% CI 88-846 days, R = −0.22, p = 0.016. ANCOVA p = 0.00018. Test for homogeneity of regressions F = 1.51, p = 0.22. **(B-E)** Immune memory at 120+ days PSO in COVID-19 non-hospitalized and hospitalized subjects. Symbol colors represent peak disease severity (white: asymptomatic, gray: mild, blue: moderate, red: severe.) For subjects with multiple sample timepoints, only the final timepoint was used for these analyses. **(B)** Spike-specific IgG (left) and RBD-specific IgG (right) binding titers. n = 64 (non-hospitalized), n = 10 (hospitalized). Mann-Whitney U tests. (**C**) Frequency memory B cells specific to Spike (left) and RBD (right) at 120+ days PSO. n = 66 (non-hospitalized), n = 10 (hospitalized). Mann-Whitney U tests. (**D**) Frequency total SARS-CoV-2-specific CD8^+^ T cells (left) and Spike-specific CD8^+^ T cells (right). p = 0.72 for total SARS-2-CoV-specific, p = 0.60 for Spike-specific by Mann-Whitney U tests. n = 72 (non-hospitalized), n = 10 (hospitalized). (**E**) Frequency total SARS-CoV-2-specific CD4^+^ T cells (left) and Spike-specific CD4^+^ T cells (right). p = 0.23 for total SARS-CoV-2-specific, p = 0.24 for Spike-specific by Mann-Whitney U tests (**F**) Immune memory to SARS-CoV-2 during the early phase (1-2 mo, black line), medium phase (3-4 mo, red line), or late phase (5-8 mo, blue line). For each individual, a score of 1 was assigned for each response above LOS for RBD IgG, Spike IgA, RBD-specific memory B cells, SARS-CoV-2 specific CD4^+^ T cells, and SARS-CoV-2-specific CD8^+^ T cells, giving a maximum total of 5 components of SARS-CoV-2 immune memory. Only COVID-19 convalescent subjects with all five immunological parameters tested were included in the analysis. n = 78 (1-2 mo), n = 52 (3-4 mo), n = 44 (5-8 mo). **(G)** Percentage dot plots showing frequencies (normalized to 100%) of subjects with indicated immune memory components as described in (B) during the early (1-2 mo) or late (5-8 mo) phase. “G”, RBD-specific IgG. “B”, RBD-specific memory B cells. “4”, SARS-CoV-2 specific CD4^+^ T cells. “8”, SARS-CoV-2 specific CD8^+^ T cells. “A”, Spike-specific IgA. n = 78 (1-2 mo), n = 44 (5-8 mo). **(H)** Relationships between immune memory compartments in COVID-19 subjects over time, as ratios (full curves and data shown in Fig. S10B-F). AU = arbitrary units, scaled from Fig. S10B-F. “B/IgA”, RBD-specific memory B cell ratio to Spike IgA antibodies. “B/IgG”, RBD-specific memory B cell ratio to RBD IgG antibodies. “B/CD4”, RBD-specific memory B cell ratio to SARS-CoV-2-specific CD4^+^ T cells. “CD4/CD8”, SARS-CoV-2-specific CD4^+^ T cells ratio to SARS-CoV-2-specific CD8^+^ T cells. “CD4/IgG”, SARS-CoV-2-specific CD4^+^ T cells ratio to RBD IgG antibodies.

Immune memory was examined for associations between magnitude of memory and COVID-19 disease severity. The number of previously hospitalized COVID-19 cases (n=13) limited analysis options. However, the cases were well distributed between males and females (**Table 1**), data from large numbers of non-hospitalized cases were available for comparison, and the analyses in Figures 1 - 4 demonstrated that immune memory was relatively stable over the time window analyzed. Therefore, we could simplify the disease severity analysis by grouping all samples from 120+ days PSO (also limiting data to a single sample per subject [Fig. S7-9]; most of the previously hospitalized subjects were sampled at two timepoints. Fig. S7A) and then comparing non-hospitalized and hospitalized subjects. Spike and RBD IgG titers in hospitalized cases were higher than non-hospitalized cases (**Fig. 5B**), consistent with other studies (*46*, *47*). Spike and RBD-specific memory B cell frequencies were also higher in hospitalized cases (~1.7-fold and ~2.5-fold, respectively. **Fig. 5C**, Fig S8). In contrast, memory CD8^+^ T cell frequencies were not higher in hospitalized cases compared to non-hospitalized cases (**Fig. 5D**, Fig. S9) and memory CD4^+^ T cell frequencies trended lower in hospitalized cases compared to non-hospitalized cases (**Fig. 5E**, Fig. S9). Therefore, while conclusions are limited by the number of hospitalized subjects, increased Spike IgG titers was consistent across three independent studies, and increased memory B cells among hospitalized cases were observed here (not measured in other studies), indicating that both compartments of long-term humoral immunity to SARS-CoV-2 are higher in individuals who experienced a more severe COVID-19 disease course. T cell memory did not follow the same pattern, consistent with indications that hospitalized cases of COVID-19 can be associated with poorer T cell responses in the acute phase (*5*, *59*). Additionally, these data show that, while gender and COVID-19 disease severity contribute to differences in immune memory to SARS-CoV-2, neither factor could account for the majority of the heterogeneity in immune memory to this virus.

Very few published data sets compare antigen-specific antibody, B cell, CD8^+^ T cell, and CD4^+^ T cell memory to an acute viral infection in the same individuals. We therefore made use of this combined data set to examine interrelationships between compartments of immune memory. We focused on RBD IgG, RBD memory B cells, Spike IgA, total SARS-CoV-2-specific CD8^+^ T cells, and total SARS-CoV-2-specific CD4^+^ T cells, due to their putative potential roles in protective immunity. The majority (64%) of COVID-19 cases were positive for all five of these immune memory compartments at 1 to 2 months PSO (**Fig. 5F-G**), with the incomplete responses largely reflecting individuals with no detectable CD8^+^ T cell memory and/or poor IgA responses (**Fig. 5G**). At 5 to 8 months after COVID-19, the proportion of individuals positive for all five of these immune memory compartments had dropped to 43%; nevertheless, 95% of individuals were still positive for at least three out of five SARS-CoV-2 immune memory responses (**Fig. 5G**). Immune memory at 5 to 8 months PSO represented contributions from different immune memory compartments in different individuals (**Fig. 5G**). Similar results were obtained if RBD IgG was replaced by neutralizing antibodies (Fig. S10A). Overall, these findings again highlight heterogeneity of immune memory, with different patterns of immune memory in different individuals.

Interrelationships between the components of memory were next examined by assessing ratios between immune memory compartments over time. The ratio of SARS-CoV-2 CD4^+^ T cell memory to SARS-CoV-2 CD8^+^ T cell memory was largely stable over time (**Fig. 5H**, Fig. S10B). Given that serological measurements are the simplest measurements of immune memory at a population scale, we examined how well such serological measurements may serve as surrogate markers of other components of SARS-CoV-2 immune memory over time. The relationship between circulating RBD IgG and RBD-specific memory B cells changed ~20-fold over the time range studied (R=0.60, **Fig. 5H,** Fig. S10C). The changing relationship between circulating Spike IgA and RBD-specific memory B cells was even larger (R=0.55, **Fig. 5H,** Fig. S10D). The relationship between RBD IgG and SARS-CoV-2 CD4^+^ T cell memory was relatively flat over the time range studied (**Fig. 5H**); however, variation spanned a ~1000-fold range (Fig. S10E). Thus, predictive power of circulating RBD IgG for assessing T cell memory was poor because of the heterogeneity between individuals (R=0.046). In sum, while heterogeneity of immune responses is a defining feature of COVID-19, immune memory to SARS-CoV-2 develops in almost all subjects, with complex relationships between the individual immune memory compartments.

## Concluding remarks

In this study, we aimed to fill gaps in our basic understanding of immune memory after COVID-19. This required simultaneous measurement of circulating antibodies, memory B cells, CD8^+^ T cells, and CD4^+^ T cells specific for SARS-CoV-2, in a group of subjects with a full range of disease, and distributed from short time points after infection out to 8 months later. By studying these multiple compartments of adaptive immunity in an integrated manner, we observed that each component of SARS-CoV-2 immune memory exhibited distinct kinetics.

The Spike IgG titers were durable, with modest declines in titers at 6 to 8 months PSO at the population level. RBD IgG and SARS-CoV-2 PSV neutralizing antibody titers were potentially similarly stable, consistent with the RBD domain of Spike being the dominant neutralizing antibody target. We collected data at two time points for most longitudinal individuals herein. It is well recognized that the magnitude of the antibody response against SARS-CoV-2 is highly heterogenous between individuals. We observed that heterogenous initial antibody responses did not collapse into a homogeneous circulating antibody memory; rather, heterogeneity is also a central feature of immune memory to this virus. For antibodies, the responses spanned a ~200-fold range. Additionally, this heterogeneity means that long-term longitudinal studies will be required to precisely define antibody kinetics to SARS-CoV-2. We are reporting the simplest statistical models that explain the data. These curve fits do not disprove more complex kinetics such as overlapping kinetics, but those models would require much denser longitudinal sampling in future studies. Nevertheless, at 5 to 8 months PSO, almost all individuals were positive for SARS-CoV-2 Spike and RBD IgG.

Notably, memory B cells specific for the Spike protein or RBD were detected in almost all COVID-19 cases, with no apparent half-life at 5 to 8 months post-infection. Other studies of RBD memory B cells are reporting similar findings (*50*, *60*). B cell memory to some other infections has been observed to be long-lived, including 60+ years after smallpox vaccination (*61*), or 90+ years after infection with influenza (*62*). The memory T cell half-lives observed over 6+ months PSO in this cohort (~125-225 days for CD8^+^ and ~94-153 days for CD4^+^ T cells) were comparable to the 123 days *t*_1/2_ observed for memory CD8^+^ T cells after yellow fever immunization (*56*). SARS-CoV-2 T cell memory at 6 months has also now been reported in another study (*63*). Notably, the durability of a fraction of the yellow fever virus-specific memory CD8^+^ T cells possessed an estimated *t*_1/2_ of 485 days by deuterium labeling (*56*). Using different approaches, the long-term durability of memory CD4^+^ T cells to smallpox, over a period of many years, was an estimated *t*_1/2_ of ~10 years (*61*, *64*), which is also consistent with recent detection of SARS-CoV-T cells 17 years after the initial infection (*65*). These data suggest that T cell memory might reach a more stable plateau, or slower decay phase, beyond the first 8 months post-infection.

While immune memory is the source of long-term protective immunity, direct conclusions about protective immunity cannot be made on the basis of quantifying SARS-CoV-2 circulating antibodies, memory B cells, CD8^+^ T cells, and CD4^+^ T cells, because mechanisms of protective immunity against SARS-CoV-2 or COVID-19 are not defined in humans. Nevertheless, some reasonable interpretations can be made. Antibodies are the only component of immune memory that can provide truly sterilizing immunity. Immunization studies in non-human primates have indicated that circulating neutralization titers of ~200 may provide sterilizing immunity against a relatively high dose URT challenge (*66*), and neutralizing titers of ~3,400 may provide sterilizing immunity against a very high dose URT challenge (*67*), although direct comparisons are not possible because the neutralizing antibody assays have not been standardized (*3*). Conclusions are also constrained by the limited overall amount of data on protective immunity to SARS-CoV-2.

Beyond sterilizing immunity, immune responses that confine SARS-CoV-2 to the URT and oral cavity would minimize COVID-19 disease severity to that of a ‘common cold’ or asymptomatic disease. This outcome is the primary goal of current COVID-19 vaccine clinical trials (*3*, *68*). Such an outcome could potentially be mediated by a mixture of memory CD4^+^ T cells, memory CD8^+^ T cells, and memory B cells specific for RBD producing anamnestic neutralizing antibodies, based on mechanisms of action in mouse models of other viral infections (*69*–*71*). In human COVID-19 infections, SARS-CoV-2-specific CD4^+^ T cells and CD8^+^ T cells are associated with less COVID-19 disease severity during an ongoing SARS-CoV-2 infection (*5*). Rapid seroconversion was associated with significantly reduced viral loads in acute disease over 14 days (*29*). Both of those associations are consistent with the hypothesis that SARS-CoV-2 memory T cells and B cells would be capable of substantially limiting SARS-CoV-2 dissemination and/or cumulative viral load, resulting in reduced COVID-19 disease severity. The likelihood of such outcomes is also closely tied to the kinetics of the infection, as memory B and T cell responses can take 3-5 days to successfully respond to an infection. As noted above, given the relatively slow course of severe COVID-19 in humans, resting immune memory compartments can potentially contribute in meaningful ways to protective immunity against pneumonia or severe secondary COVID-19. The presence of sub-sterilizing neutralizing antibody titers at the time of SARS-CoV-2 exposure would blunt the size of the initial infection, and may provide an added contribution to limiting COVID-19 severity, based on observations of protective immunity for other human respiratory viral infections (*37*, *72*–*74*) and observations of SARS-CoV-2 vaccines in non-human primates (*48*, *67*, *75*).

The current study has some limitations. Longitudinal data for each subject, with at least three time points per subject, would be required for more precise understanding of the kinetics of durability of SARS-CoV-2 antibodies. Nevertheless, the current cross-sectional data describe well the dynamics of SARS-CoV-2 memory B cells, CD8^+^ T cell, and CD4^+^ T cell over 8 months PSO. This study was not sufficiently powered to control for many variables simultaneously. Additionally, circulating memory was assessed here; it is possible that local URT immune memory is a minimal, moderate, or large component of immune memory after a primary infection with SARS-CoV-2. This remains to be determined.

Individual case reports show that reinfections with SARS-CoV-2 are occurring (*76*, *77*). However, a 2,800 person study found no symptomatic re-infections over a ~118 day window (*78*), and a 1,246 person study observed no symptomatic re-infections over 6 months (*79*). We observed heterogeneity in the magnitude of adaptive immune responses to SARS-CoV-2 persisting into the immune memory phase. It is therefore possible that a fraction of the SARS-CoV-2-infected population with low immune memory would become susceptible to re-infection relatively soon. While gender and disease severity both contribute some to the heterogeneity of immune memory reported here, the source of much of the heterogeneity in immune memory to SARS-CoV-2 is unknown and worth further examination. Perhaps heterogeneity derives from low cumulative viral load or a small initial inoculum in some individuals. Nevertheless, our data show immune memory in at least three immunological compartments was measurable in ~95% of subjects 5 to 8 months PSO, indicating that durable immunity against secondary COVID-19 disease is a possibility in most individuals.

## ACKNOWLEDGEMENTS

We would like to thank the LJI Clinical Core, specifically Gina Levi, RN and Brittany Schwan for healthy donor enrollment and blood sample procurement. We thank Carolyn Moderbacher for input on data analysis. We are also grateful to the Mt. Sinai Personalized Virology Initiative for sharing banked samples from study participants with COVID-19. We are grateful to Dr. A. Wajnberg for study participant referrals and to the Personalized Virology Initiative (Dr. G. Kleiner, Dr. LCF Mulder, Dr. M. Saksena, K. Srivastava, C. Gleason, C. M. Bermúdez-González, K. Beach, K. Russo, L. Sominsky, E. Ferreri, R. Chernet, L. Eaker, A. Salimbangon, D. Jurczyszak, H. Alshammary, W. Mendez, A. Amoako, S. Fabre, S. Suthakaran, M. Awawda, E. Hirsch, A. Shin) for sharing banked samples from study participants with COVID-19.

## Authors contributions

Conceptualization, S.C., A.S. and D.W.; Investigation, J.M.D., J.M., Y.K., K.M.H., E.D.Y., C.E.F., A.G., S.H., C.N.; Formal Analysis, J.M.D., J.M., Y.K., K.M.H., C.E.F., S.H., B.P., D.W., A.S., S.C.; Patient Recruitment and Samples, S.I.R., A.F., S.A.R., F. K., V. S., D.M.S., D.W.; Material Resources, F.K., V.S., V.R., E.O.S., D.W., A.S., S.C.; Data Curation, Y.K., J.M.D., J.M., S.H.; Writing, Y.K., J.M.D., J.M., S.I.R., D.W., A.S., S.C.; Supervision, D.W., A.S., S.C., Project Administration, A.F.

## Data and materials availability

All data are provided in the Supplementary Materials. Epitope pools utilized in this paper will be made available to the scientific community upon request and execution of a material transfer agreement (MTA). This work is licensed under a Creative Commons Attribution 4.0 International (CC BY 4.0) license, which permits unrestricted use, distribution, and reproduction in any medium, provided the original work is properly cited. To view a copy of this license, visit https://creativecommons.org/licenses/by/4.0/. This license does not apply to figures/photos/artwork or other content included in the article that is credited to a third party; obtain authorization from the rights holder before using such material.

## Supplementary Materials

Materials and Methods

Figures S1-S10

Tables S1-S2

Data File

## Supplementary Materials for

### Materials and Methods

#### Human Subjects

The Institutional Review Boards of the University of California, San Diego (UCSD; 200236X) and the La Jolla Institute for Immunology (LJI; VD-214) approved the protocols used for blood collection for subjects with COVID-19 who donated at all sites other than Mt. Sinai. The Icahn School of Medicine at Mt. Sinai IRB approved the samples collected at this institution in New York City (IRB-16-00791). All human subjects were assessed for medical decision-making capacity using a standardized, approved assessment, and voluntarily gave informed consent prior to being enrolled in the study. Study inclusion criteria included a diagnosis of COVID-19 or suspected COVID-19, age of 18 years or greater, willingness and ability to provide informed consent. Although not a strict inclusion criterion, evidence of positive PCR-based testing for SARS-CoV-2 was requested from subjects prior to participation. 145 cases were confirmed SARS-CoV-2 positive by PCR-based testing (**Table 1**). Two subjects tested negative by SARS-CoV-2 PCR (**Table 1**). The remainder were not tested or did not have test results available for review (**Table 1**). Subjects who had a medical history and/or symptoms consistent with COVID-19, but lacked positive PCR-based testing for SARS-CoV-2 and subsequently had negative laboratory-based serologic testing for SARS-CoV-2 were then excluded; i.e., all COVID-19 cases in this study were confirmed cases by SARS-CoV-2 PCR or SARS-CoV-2 serodiagnostics, or both. Adults of all races, ethnicities, ages, and genders were eligible to participate. Study exclusion criteria included lack of willingness to participate, lack of ability to provide informed consent, or a medical contraindication to blood donation (e.g. severe anemia). Subject samples at LJI were obtained from individuals in California and at least seven other states.

Blood collection and processing methods at LJI were performed as previously described (*5*). Briefly, whole blood was collected via phlebotomy in acid citrate dextrose (ACD) serum separator tubes (SST), or ethylenediaminetetraacetic acid (EDTA) tubes and processed for peripheral blood mononuclear cells (PBMC), serum, and plasma isolation. Most donors were screened for symptoms prior to scheduling blood draws, and had to be symptom-free and approximately 3-4 weeks out from symptom onset at the time of the initial blood draw at UCSD or LJI, respectively. Samples were coded, and then de-identified prior to analysis. Other efforts to maintain the confidentiality of participants included the labeling samples with coded identification numbers. An overview of the characteristics of subjects with COVID-19 is provided in **Table 1**.

COVID-19 disease severity was scored from 0 to 10 using a numerical scoring system based on the NIH ordinal scale (*5*, *80*). A categorical descriptor was applied based on this scoring system: “asymptomatic” for a score of 1, “mild” for a score of 2-3, “moderate” for a score of 4-5, and “severe” for a score of 6 or more. Subjects with a numerical score of 4 or higher required hospitalization (including admission for observation) for management of COVID-19. Only one of 13 hospitalized subjects is shared from the previous study of acute COVID-19 (*5*). The days PSO was determined based on the difference between the date of the blood collection and the date of first reported symptoms consistent with COVID-19. For asymptomatic subjects, the day from first positive SARS-CoV-2 PCR-based testing was used in place of the date of first reported COVID-19 symptoms.

#### Recombinant Proteins

Stabilized Spike protein (2P, (*81*)) and the receptor binding domain (RBD) were expressed in HEK293F cells. Briefly, DNA expressing stabilized spike protein and RBD were subcloned into separate phCMV vectors and transfected into HEK293F cells at a ratio of 1mg of DNA to 1L of cells. The cells were cultured at 37C in a shaker incubator set to 125rpm, 80% humidity and 8% CO2. When cell viability dropped below 80% (typically 4-5 days), media was harvested and centrifuged to remove cells. Biolock reagent was added to the supernatant media to remove any excess biotin. The media was then filtered through a 0.22um filter to remove Biolocked-aggregates. Proteins were purified using Streptrap HP 5mL columns (Cytiva) using 100mM Tris, 100mM NaCl as the Wash Buffer and 100mM Tris, 100mM NaCl, 2.5mM d-Desthiobiotin as the Elution Buffer. The eluted fractions for Spike proteins were concentrated on 100kDa Amicon filters while the RBD were concentrated on 10kDa filters. The samples were further purified using S6increase columns for the spike variants and S200increase column for RBD.

#### SARS-CoV-2 ELISAs

SARS-CoV-2 ELISAs were performed as previously described (*2, 5, 82).* Briefly, Corning 96-well half area plates (ThermoFisher 3690) were coated with 1μg/mL of antigen overnight at 4°C. Antigens included recombinant SARS-CoV-2 RBD protein, recombinant Spike protein, and recombinant Nucleocapsid protein (GenScript Z03488) (Recombinant nucleocapsid antigens were also tested from Sino Biological (40588-V07E) and Invivogen (his-sars2-n) and yielded comparable results to GenScript nucleocapsid). The following day, plates were blocked with 3% milk in phosphate buffered saline (PBS) containing 0.05% Tween-20 for 1.5 hours at room temperature. Plasma was heat inactivated at 56°C for 30-60 minutes. Plasma was diluted in 1% milk containing 0.05% Tween-20 in PBS starting at a 1:3 dilution followed by serial dilutions by 3 and incubated for 1.5 hours at room temperature. Plates were washed 5 times with 0.05% PBS-Tween-20. Secondary antibodies were diluted in 1% milk containing 0.05% Tween-20 in PBS. For IgG, anti-human IgG peroxidase antibody produced in goat (Sigma A6029) was used at a 1:5,000 dilution. For IgA, anti-human IgA horseradish peroxidase antibody (Hybridoma Reagent Laboratory HP6123-HRP) was used at a 1:1,000 dilution. The HP6123 monoclonal anti-IgA was used because of its CDC and WHO validated specificity for human IgA1 and IgA2 and lack of crossreactivity with non-IgA isotypes (*82*).

Endpoint titers were plotted for each sample, using background subtracted data. Negative and positive controls were used to standardize each assay and normalize across experiments. A positive control standard was created by pooling plasma from 6 convalescent COVID-19 donors to normalize between experiments. The limit of detection (LOD) was defined as 1:3 for IgG, 1:10 for IgA. Limit of sensitivity (LOS) for SARS-CoV-2 infected individuals was established based on uninfected subjects, using plasma from normal healthy donors never exposed to SARS-CoV-2. For cross-sectional analyses, modeling for the best fit curve (e.g., one phase decay versus simple linear regression) was performed using GraphPad Prism 8.0. Best curve fit was defined by an extra sum-of-squares F Test, selecting the simpler model unless P < 0.05 (*83).* Continuous decay (linear regression), one-phased decay, or two-phased decay of log data were assessed in all cases, with the best fitting statistical model chosen based on the F test; in several cases a quadratic equation fit was also considered. To calculate the *t*_1/2_, log2 transformed data was utilized. Using the best fit curve, either a one phase decay non-linear fit or a simple linear regression (continuous decay) was utilized. For simple linear regressions, Pearson R was calculated for correlation using log2 transformed data. For one phase decay non-linear fit, R was reported. For longitudinal samples, a simple linear regression was performed, with *t1/2* calculated from log2 transformed data for each pair. For gender analyses, modeling and *t*_1/2_ was performed similar to cross-sectional analyses; ANCOVA (VassarStats or GraphPad Prism 8.4) was then performed between male and female data sets. ANCOVA p-values of the adjusted means were reported and considered significant if the test for homogeneity of regressions was not significant.

#### Neutralizing antibody assays

The pseudovirus neutralizing antibody assay was performed as previously described (*5*). Briefly, Vero cells were seeded in 96-well plates to produce a monolayer at the time of infection. Pre-titrated amounts of rVSV-SARS-Cov-2 (phCMV3-SARS-CoV-2 Spike SARS-CoV-2-pseduotyped VSV-ΔG-GFP were generated by transfecting HEK293T cells, ATCC CRL-3216) were incubated with serially diluted human plasma at 37°C for 1 hour before addition to confluent Vero cell monolayers (ATCC CCL-81) in 96-well plates. Cells were incubated for 12-16 hours at 37°C in 5% CO2. Cells were then fixed in 4% paraformaldehyde, stained with 1μg/mL Hoechst, and imaged using a CellInsight CX5 imager to quantify the total number of cells expressing GFP. Infection was normalized to the average number of cells infected with rVSV-SARS-CoV-2 incubated with normal human plasma. The limit of detection (LOD) was established as < 1:20 based on plasma samples from a series of unexposed control subjects. Negative signals were set to 1:19. Neutralization IC50 titers were calculated using One-Site Fit LogIC50 regression in GraphPad Prism 8.0.

#### Detection of antigen-specific memory B cells

To detect SARS-CoV-2 specific B cells, biotinylated protein antigens were individually multimerized with fluorescently labeled streptavidin at 4°C for one hour. Full-length SARS-CoV-2 Spike (2P-stabilized, double Strep-tagged) and RBD were generated in-house. Biotinylation was performed using biotin-protein ligase standard reaction kit (Avidity, Cat# Bir500A) following the manufacturer’s standard protocol and dialyzed overnight against PBS. Biotinylated Spike was mixed with streptavidin BV421 (BioLegend, Cat# 405225) and streptavidin Alexa Fluor 647 (Thermo Fisher Scientific, Cat# S21374) at 20:1 ratio (~6:1 molar ratio). Biotinylated RBD was mixed with streptavidin PE/Cyanine7 (BioLegend, Cat# 405206) at 2.2:1 ratio (~4:1 molar ratio). Biotinylated SARS-CoV-2 full length Nucleocapsid (Avi- and His-tagged; Sino Biological, Cat# 40588-V27B-B) was multimerized using streptavidin PE (BioLegend, Cat# 405204) and streptavidin BV711 (BioLegend, Cat# 405241) at 5.5:1 ratio (~6:1 molar ratio). Streptavidin PE/Cyanine5.5 (Thermo Fisher Scientific, Cat# SA1018) was used as a decoy probe to gate out SARS-CoV-2 non-specific streptavidin-binding B cells. The antigen probes prepared individually as above were then mixed in Brilliant Buffer (BD Bioscience, Cat# 566349) containing 5μM free d-biotin (Avidity, Cat# Bir500A). Free d-biotin ensured minimal cross-reactivity of antigen probes. ~10^7^ previously frozen PBMC samples were prepared in U-bottom 96-well plates and stained with 50μL antigen probe cocktail containing 100ng Spike per probe (total 200ng), 27.5ng RBD, 40ng Nucleocapsid per probe (total 80ng) and 20ng streptavidin PE/Cyanine5.5 at 4°C for one hour to ensure maximal staining quality before surface staining with antibodies as listed in Table S1 was performed in Brilliant Buffer at 4°C for 30min. Dead cells were stained using LIVE/DEAD Fixable Blue Stain Kit (Thermo Fisher Scientific, Cat# L34962) in DPBS at 4°C for 30min. ~80% of antigen-specific memory (IgD^-^ and/or CD27^+^) B cells detected using this method were IgM^+^, IgG^+^, or IgM^-^ IgG IgA^+^, which were comparable to non-specific memory B cells. Based on these observations, we concluded that the antigen probes did not significantly impact the quality of surface immunoglobulin staining. Stained PBMC samples were acquired on Cytek Aurora and analyzed using FlowJo10.7.1 (BD Bioscience).

The frequency of antigen-specific memory B cells was expressed as a percentage of total B cells (CD19^+^ CD20^+^ CD38^int/-^, CD3^-^, CD14^-^, CD16^-^, CD56^-^, LIVE/DEAD^-^, lymphocytes), or as number per 10^6^ PBMC (LIVE/DEAD^-^ cells). LOD was set based on median + 2× standard deviation (SD) of [1 / (number of total B cells recorded)] or median + 2×SD of [10^6^ / (number of PBMC recorded)]. LOS was set as the median + 2×SD of the results in unexposed donors. Phenotype analysis of antigen-specific B cells was performed only in subjects with at least 10 cells detected in the respective antigen-specific memory B cell gate. In each experiment, PBMC from a known positive control (COVID-19 convalescent subject) and unexposed subjects were included to ensure consistent sensitivity and specificity of the assay. For each data set, second order polynomial, simple linear regression, and pseudo-first order kinetic models were considered. The model with a lower Akaike’s Information Criterion value was determined to be a better-fit and visualized.

#### Activation induced markers (AIM) T cell assay

Antigen-specific CD4^+^ T cells were measured as a percentage of AIM^+^ (OX40^+^CD137^+^) CD4^+^ T and (CD69^+^CD137^+^) CD8^+^ T cells after stimulation of PBMC with overlapping peptide megapools (MP) spanning the entire SARS-CoV-2 ORFeome, as previously described (*2*). Cells were cultured for 24 hours in the presence of SARS-CoV-2 specific MPs [1 μg/mL] or 5 μg/mL phytohemagglutinin (PHA, Roche) in 96-wells U-bottom plates at 1×10^6^ PBMC per well. A stimulation with an equimolar amount of DMSO was performed as a negative control, PHA, and stimulation with a combined CD4^+^ and CD8^+^ cytomegalovirus epitope MP (CMV, 1 μg/mL) were included as positive controls. Any sample with low PHA signal was excluded as a quality control.

Antigen-specific CD4^+^ and CD8^+^ T cells were measured as background (DMSO) subtracted data, with a minimal DMSO level set to 0.005%. All positive ORFs (> 0.02% for CD4^+^, > 0.05% for CD8^+^) were then aggregated into a combined sum of SARS-CoV-2-specific CD4^+^ or CD8^+^ T cells. The threshold for positivity for antigen-specific CD4^+^ T cell responses (0.03%) and antigen-specific CD8^+^ T cell responses (0.12%) was calculated using the median two-fold standard deviation of all negative controls measured (>150). The antibody panel utilized in the (OX40^+^CD137^+^) CD4^+^ T and (CD69^+^CD137^+^) CD8^+^ T cells AIM staining is shown in Table S2. A consistency analysis was performed for multiple measurements of AIM T cell assays by two different operators. Before merging, we compared the protein immunodominance, total SARS-CoV-2-specific CD4^+^ and CD8^+^ T cell responses, and half-life calculations between the two groups of experimental data. In longitudinal analyses, half-life calculations excluded any samples that were negative at both timepoints (since a half-life could not be calculated), though all data were included in the graphs.

For surface CD40L^+^OX40^+^ CD4^+^ T cell AIM assays, experiments were performed as previously described (*5*), with the following modifications. Cells were cultured in complete RPMI containing 5% human AB serum (Gemini Bioproducts), beta-mercaptoethanol, Penicillin/Streptomycin, sodium pyruvate (NaPy), and non-essential amino acids. Prior to addition of peptide MPs, cells were blocked at 37°C for 15 minutes with 0.5μg/mL anti-CD40 mAb (Miltenyi Biotec). A stimulation with an equimolar amount of DMSO was performed to determine background subtraction, and activation from Staphylococcal enterotoxin B (SEB) at 1 μg/mL was used as (positive) quality control. LOD for antigen-specific cTFH among CD4^+^ T cells was based on the LOD for antigen-specific CD4^+^ T cells (described above) multiplied by the average % cTFH in the bulk CD4 T cells among control samples. An inclusion threshold of ten events after the cTFH CXCR5^+^ gate was used for PD-1^hi^ and CCR6^+^ calculations, and Mann-Whitney nonparametric and Wilcoxon signed-rank statistical tests were applied for the respective comparisons.

---

**Fig. S1.**
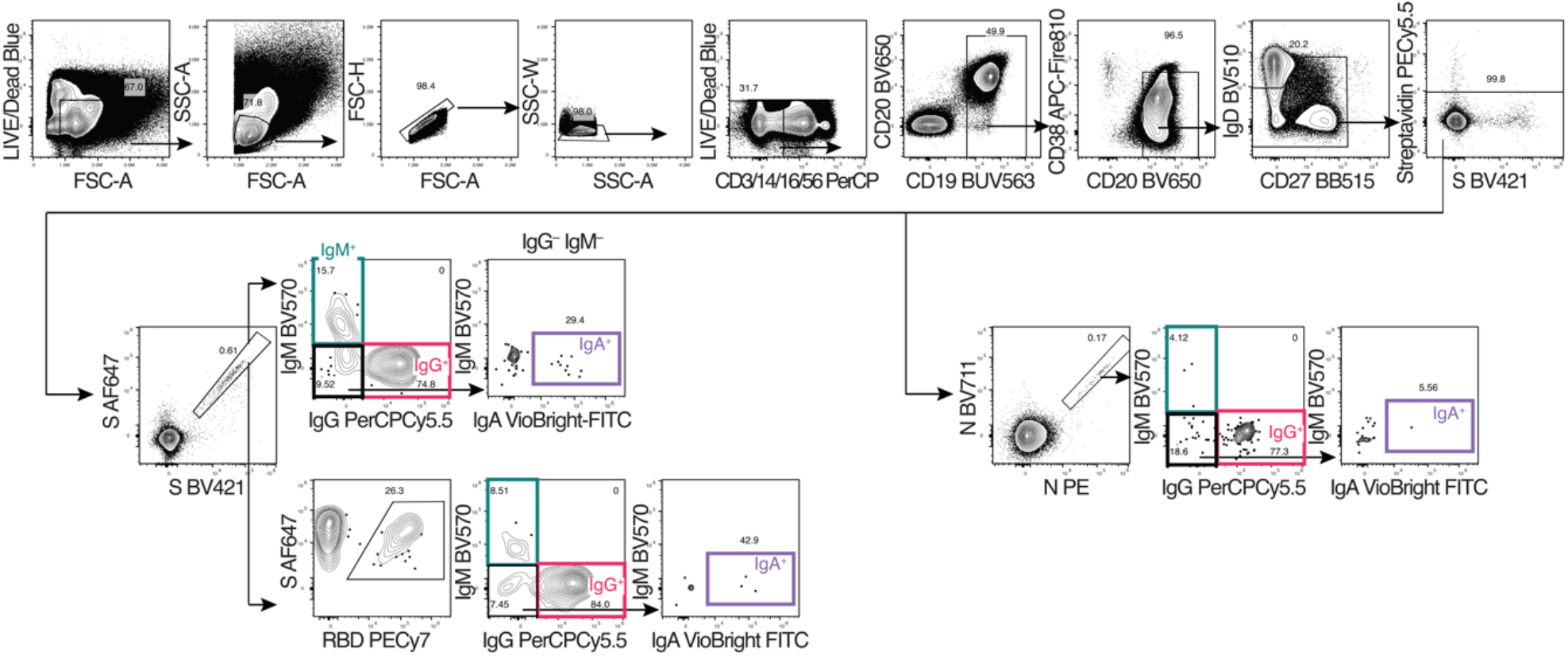
SARS-CoV-2 memory B cells. (**A**) Gating strategies to define Spike-, RBD-, or Nucleocapsid-specific memory B cells. S = SARS-CoV-2 Spike trimer. N = SARS-CoV-2 Nucleocapsid.

**Fig. S2.**
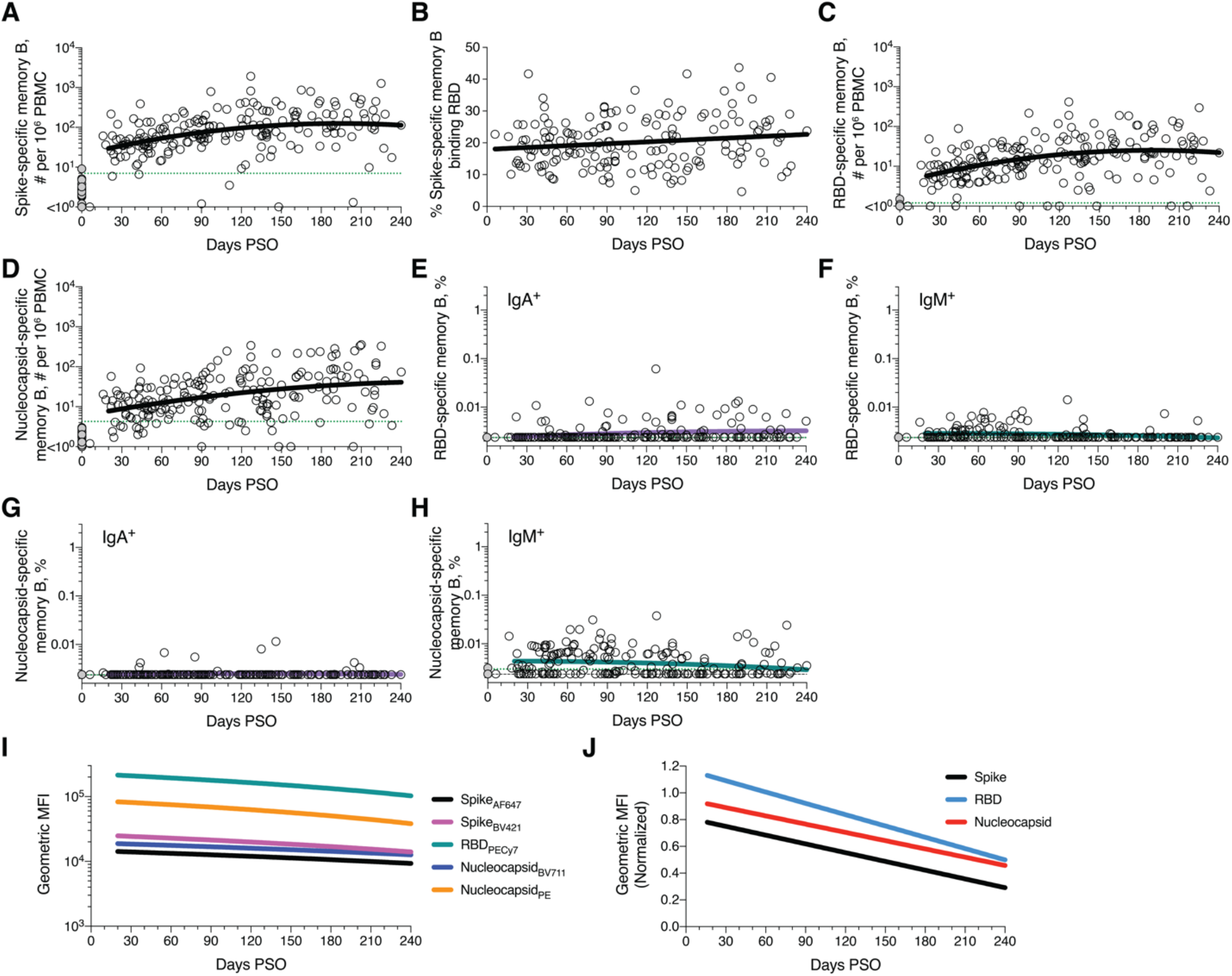
Kinetics of memory B cell responses. (**A**) Cross-sectional analysis of SARS-CoV-2 Spike-specific memory B cell numbers per 10^6^ PBMC. Second order polynomial model for best fit curve (|R| = 0.38). **(B)** Percentage of Spike-specific B cells binding RBD. Simple linear regression (R = 0.15) (**C**) Cross-sectional analysis of RBD-specific memory B cell numbers per 10^6^ PBMC. Second order polynomial model for best fit curve (|R| = 0.39). (**D**) Cross-sectional analysis of Nucleocapsid-specific memory B cell numbers per 10^6^ PBMC. Second order polynomial model for best fit curve (|R| = 0.38). (**E**) Cross-sectional analysis of frequency (% of CD19^+^ CD20^+^ B cells) of RBD-specific IgA^+^ memory B cells. Second order polynomial model for best fit curve (|R| = 0.19). (**F**) Cross-sectional analysis of frequency of RBD-specific IgM^+^ memory B cells. Second order polynomial model for best fit curve (|R| = 0.18). (**G**) Cross-sectional analysis of frequency of SARS-CoV-2 Nucleocapsid-specific IgA^+^ memory B cells. Second order polynomial model (|R| = 0.06). (**H**) Cross-sectional analysis of frequency of Nucleocapsid-specific IgM^+^ memory B cells. Second order polynomial (|R| = 0.17). (**I**) Cross-sectional analysis of geometric mean fluorescence intensity of Spike, RBD and Nucleocapsid probes on S-, RBD- and Nucleocapsid-specific memory B cells, respectively. Data shown are simple linear-regression lines. (**J**) Cross-sectional analysis of geometric mean fluorescence intensity of Spike, RBD and Nucleocapsid probes on S-, RBD- and Nucleocapsid-specific memory B cells, respectively, normalized to a positive control sample. Data shown are simple linear-regression lines.

**Fig. S3.**
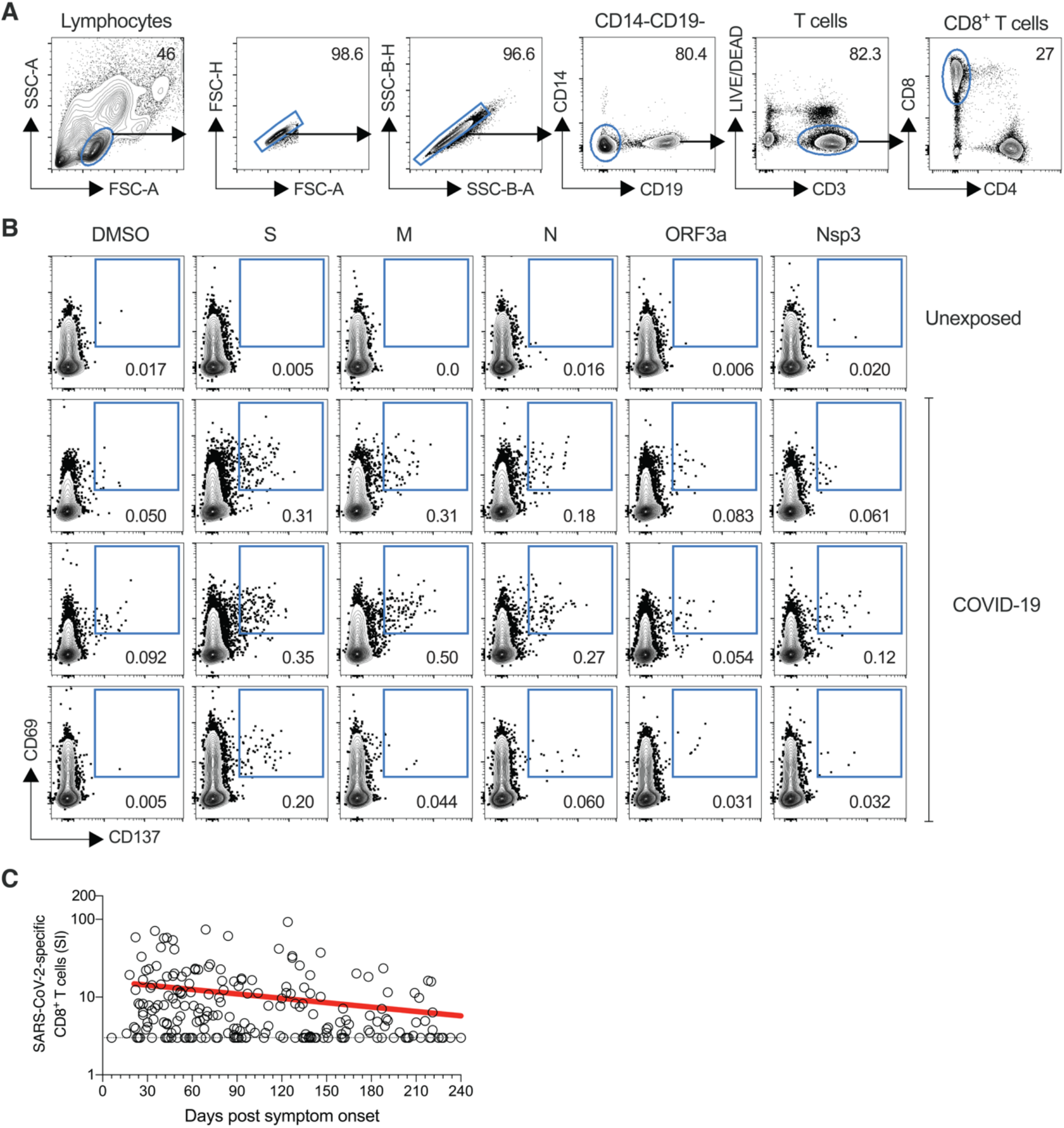
SARS-CoV-2 circulating memory CD8^+^ T cells. **(A)** Gating strategies to define SARS-CoV-2-specific CD8^+^ T cells by AIM assay, using individual SARS-CoV-2 ORF peptide pools. **(B)** Representative examples of flow cytometry plots of SARS-CoV-2-specific CD8^+^ T cells (CD69^+^ CD137^+^) after overnight stimulation with Spike (S), Membrane (M), Nucleocapsid (N), ORF3a, or nsp3 peptide pools, compared to negative control stimulation (DMSO) from three COVID-19 subjects and one uninfected control. **(C)** Crosssectional analysis of total SARS-CoV-2-specific CD8^+^ T cells, as per Figure 3, but graphing stimulation index (SI).

**Fig. S4.**
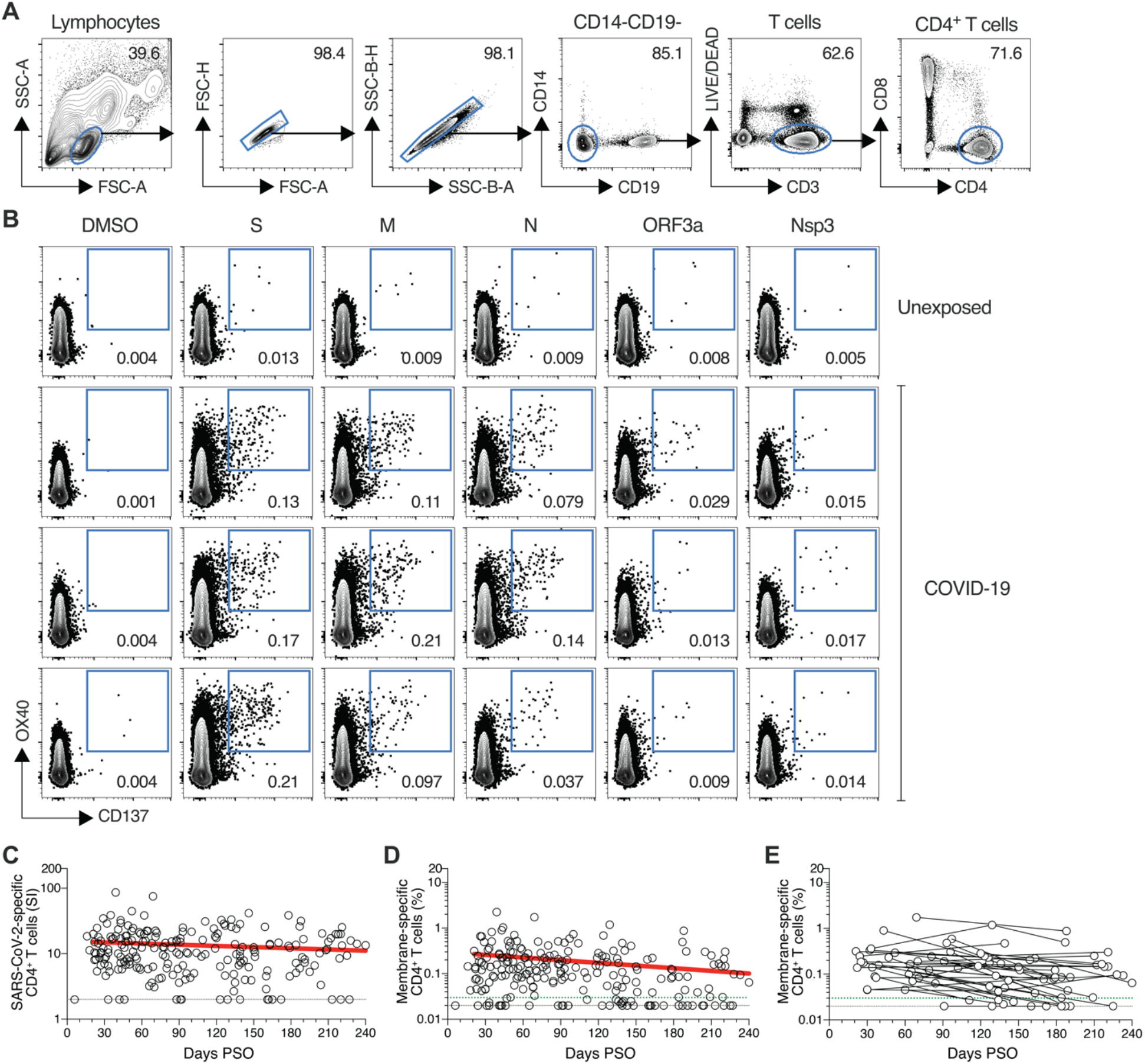
SARS-CoV-2 circulating memory CD4^+^ T cells. **(A)** Gating strategies to define SARS-CoV-2-specific CD4^+^ T cells by AIM assay, using individual SARS-CoV-2 ORF peptide pools. (**B)** Representative examples of flow cytometry plots of SARS-CoV-2-specific CD4^+^ T cells (OX40^+^ CD137^+^, after overnight stimulation with Spike, M, Nucleocapsid, ORF3a, or nsp3 peptide pools, compared to negative control (DMSO). From three COVID-19 subjects and one uninfected control. **(C)** Cross-sectional analysis of total SARS-CoV-2-specific CD4^+^ T cells, as per Figure 4, but graphing stimulation index (SI). **(D)** Cross-sectional analysis of M-specific CD4^+^ T cells. Linear decay preferred model, *t*_1/2_ = 153 days. R = −0.25, p = 0.0003. **(E)** Longitudinal analysis of M-specific CD4^+^ T cells in paired samples from the same subjects. n = 215 COVID-19 subject samples for cross-sectional analysis. n = 37 COVID-19 subjects for longitudinal analysis.

**Fig. S5.**
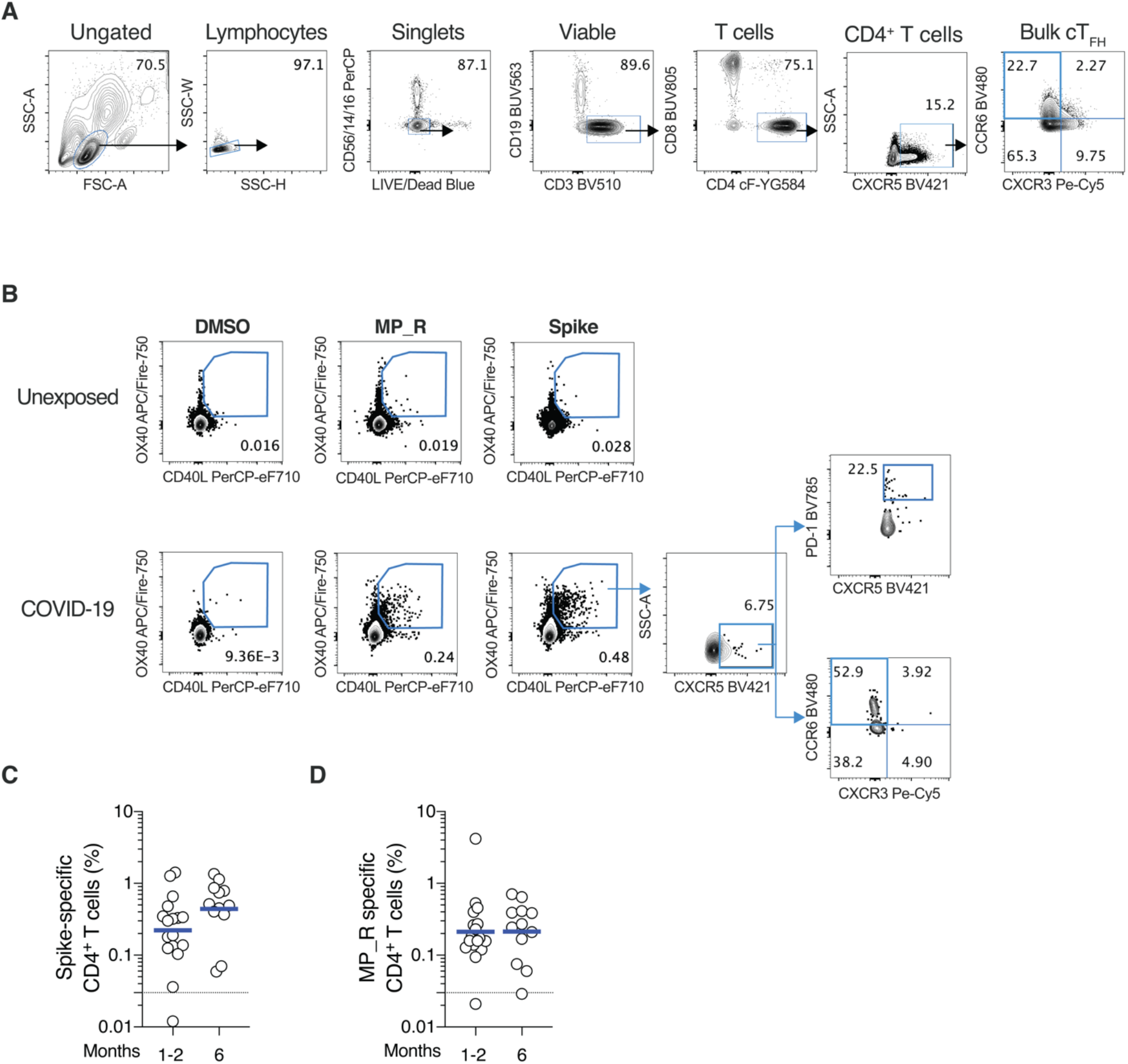
SARS-CoV-2 memory T_FH_ cells. **(A)** Gating strategies to define SARS-CoV-2-specific CD4^+^ T cells by AIM assay, using Spike and MP_R peptide pools. **(B)** Representative examples of flow cytometry plots of SARS-CoV-2-specific CD4^+^ T cells. Surface CD40L^+^OX40^+^, after overnight stimulation with Spike or MP_R peptide pools, compared to negative control (DMSO) from a representative COVID-19 subject and an uninfected control. **(C, D)** SARS-CoV-2-specific CD4^+^ T cells based on surface CD40L^+^OX40^+^, gated as in A, after overnight stimulation with Spike or MP_R peptide pools. n = 29 COVID-19 subject samples (white circles), n = 17 at 1-2 mo, n = 12 at 6 mo.

**Fig. S6.**
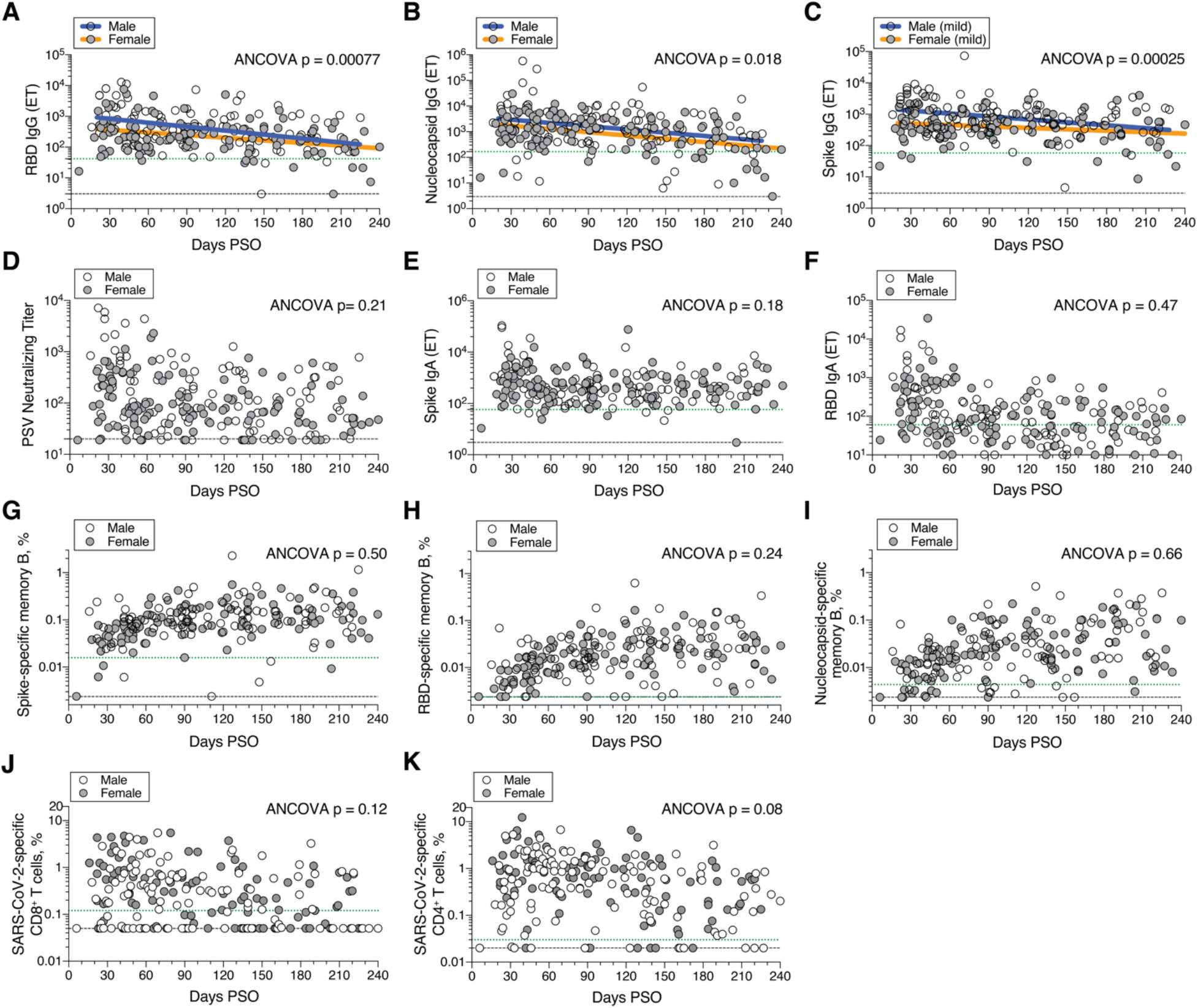
Immune memory and gender. Cross-sectional analyses of SARS-CoV-2 immune memory by male and female gender. **(A)** RBD IgG titers. Males: Continuous decay preferred model, initial *t*_1/2_ = 24 days, R = −0.39, p<0.0001. Females: linear decay preferred model, *t*_1/2_ = 94 days, 95% CI: 64-178 days R = −0.36, p<0.0001. ANCOVA = 0.00077, Test for homogeneity of regressions F = 1.32, p = 0.25. **(B)** Nucleocapsid IgG titers. Males: Continuous decay preferred model, *t*_1/2_ = 70 days, 95% CI: 42-209 days, R = −0.28, p = 0.0035. Females: continuous decay preferred model, *t*_1/2_ = 64 days, 95% CI: 47-104 days, R = −0.44, p < 0.0001. ANCOVA p = 0.018, Test for homogeneity of regressions F = 0, p = 1.0. **(C)** Spike IgG titers of non-hospitalized patients. Males: Continuous decay preferred model *t*_1/2_ = 574 days, 95% CI: 345-1698 days, R = −0.30, p = 0.0035. Females: continuous decay preferred model, *t*_1/2_ = 1075 days, 95% CI: 537-1,340,303 days, R = −0.19, p = 0.0502. ANCOVA p = 0.00025, Test for homogeneity of regressions F = 2.59, p = 0.11. **(D)** PSV neutralizing titers. **(E)** Spike IgA titers. **(F)** RBD IgA titers. **(G)** Frequency (% of CD19^+^ CD20^+^ B cells) of SARS-CoV-2 Spike-specific total (IgG^+^, IgA^+^, or IgM^+^) memory B cells, as per Figure 2C. **(H)** Frequency of SARS-CoV-2 RBD-specific total memory B cells, as per Figure 2E. **(I)** Frequency of SARS-CoV-2 Nucleocapsid-specific total memory B cells, as per Figure 2G. **(J)**. Frequency (% of CD8^+^ T cells) of total SARS-CoV-2-specific CD8^+^ T cells, as per Figure 3B. **(K)** Frequency (% of CD4^+^ T cells) of total SARS-CoV-2-specific CD4^+^ T cells, as per Figure 4B.

**Fig. S7.**
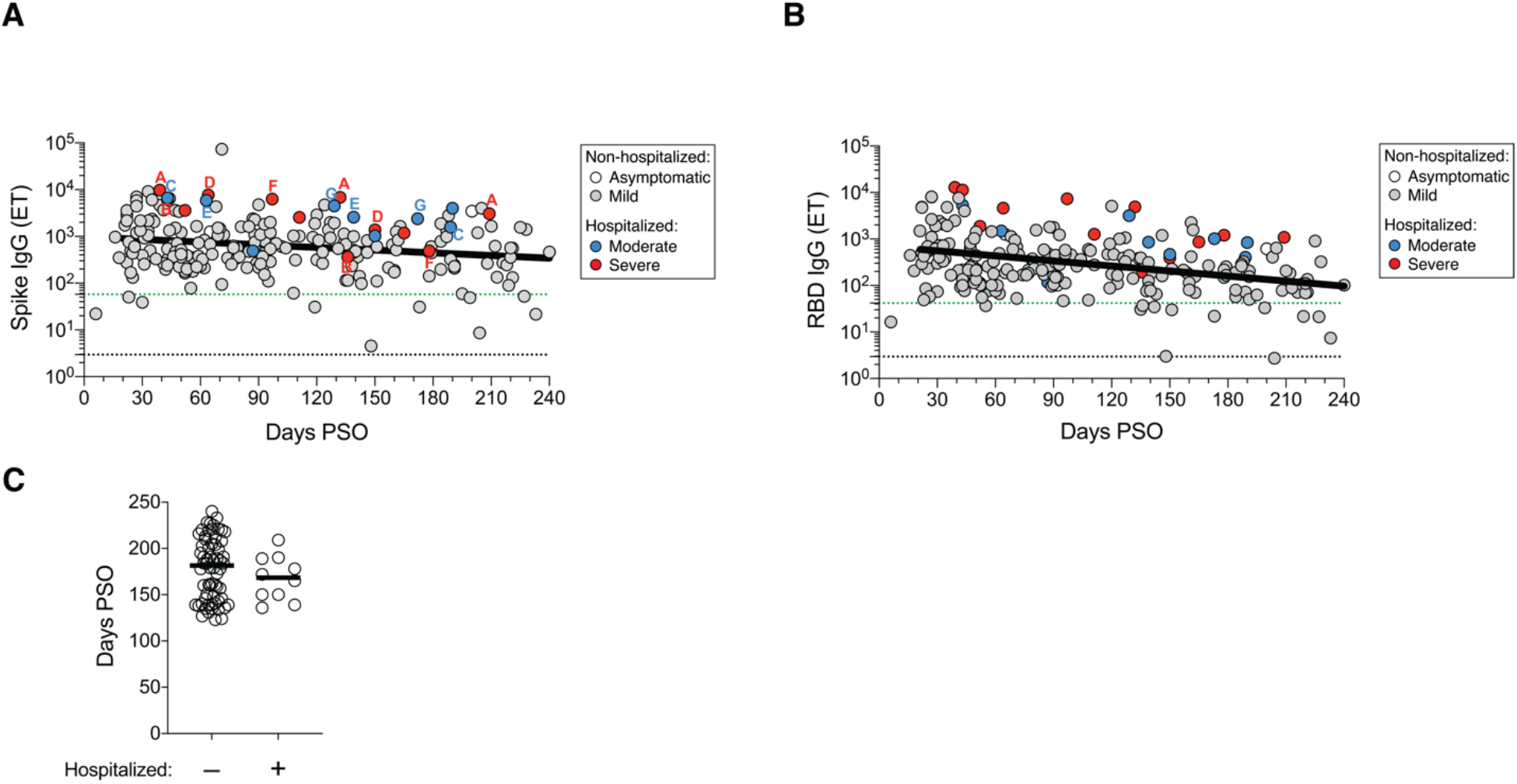
Serological memory and disease severity. **(A)** Cross-sectional analysis of Spike IgG, as per Figure 1A, color coded based on subject COVID-19 disease severity (white: asymptomatic, gray: mild, blue: moderate, red: severe). Letters indicate donors that were sampled at multiple timepoints after the onset of symptoms. One letter per donor. **(B)** Cross-sectional analysis of RBD IgG, as per Figure 1C, color coded based on subject COVID-19 disease severity. (C) Distribution of timepoints of COVID-19 convalescent subjects (120+ days PSO) analyzed in Figure 5B. Line indicates median. For subjects with multiple sample timepoints, only the final timepoint was used for these analyses. *p* = 0.40, Mann-Whitney test.

**Fig. S8.**
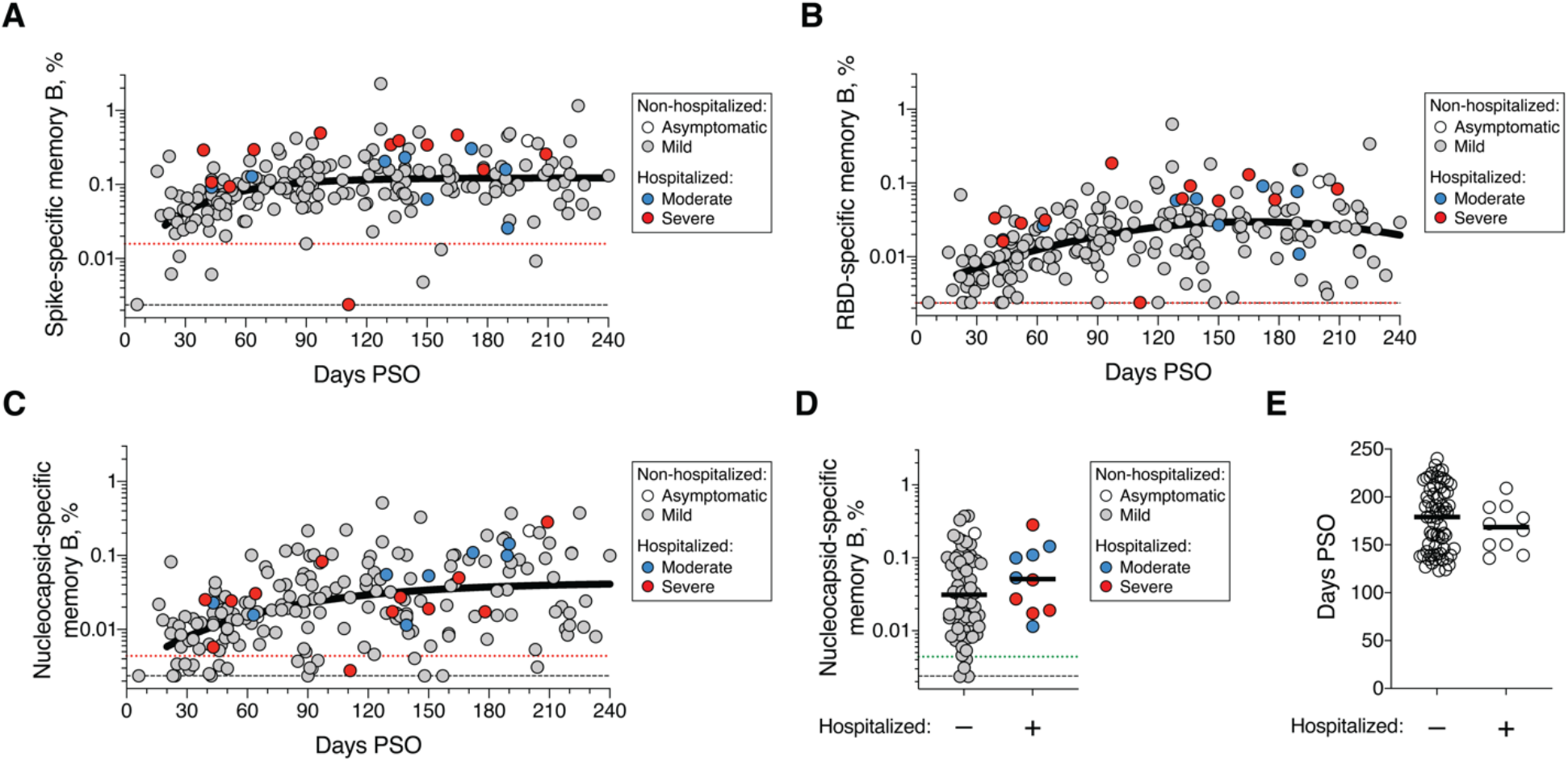
Memory B cells and disease severity. **(A)** Cross-sectional analysis of SARS-CoV-2 Spike-specific total (IgG^+^, IgA^+^, or IgA^+^) memory B cells, as per Figure 2C, color coded based on subject COVID-19 disease severity (white: asymptomatic, gray: mild, blue: moderate, red: severe). **(B)** Cross-sectional analysis of RBD-specific total memory B cells, as per Figure 2E, color coded based on subject COVID-19 disease severity. **(C)** Cross-sectional analysis of Nucleocapsid-specific total memory B cells, as per Figure 2G, color coded based on subject COVID-19 disease severity. (**D**) Frequency of Nucleocapsid-specific memory B cells at 120+ days PSO in non-hospitalized (Asymptomatic and Mild) and hospitalized cases (Moderate and Severe). *p* = 0.20, Mann-Whitney test. **(E)** Distribution of timepoints of COVID-19 convalescent subjects (120+ days PSO) analyzed in Figure 5C, S8D. Line indicates median. For subjects with multiple sample timepoints, only the final timepoint was used for these analyses. *p* = 0.47, Mann-Whitney test.

**Fig. S9.**
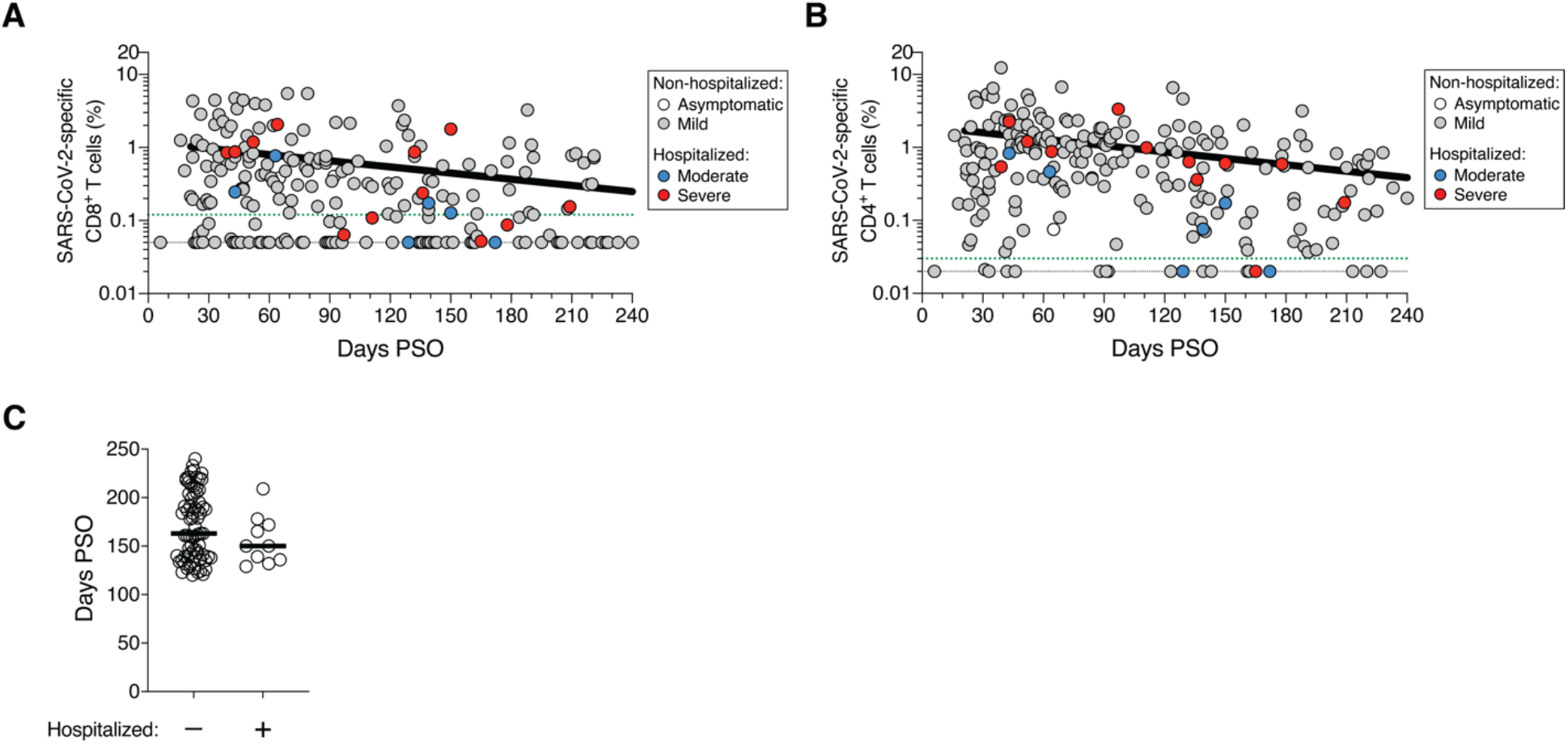
T cell memory and disease severity. **(A)** Cross-sectional analysis of SARS-CoV-2-specific CD8^+^ T cells, as per Figure 3B, color coded based on subject COVID-19 disease severity (white: asymptomatic, gray: mild, blue: moderate, red: severe). **(B)** Cross-sectional analysis of SARS-CoV-2-specific CD4^+^ T cells, as per Figure 4B, color coded based on subject COVID-19 disease severity. **(C)** Distribution of timepoints of COVID-19 convalescent subjects (120+ days PSO) analyzed in Figure 5D-E. Line indicates median. For subjects with multiple sample timepoints, only the final timepoint was used for these analyses. *p* = 0.23, Mann-Whitney test.

**Fig. S10.**
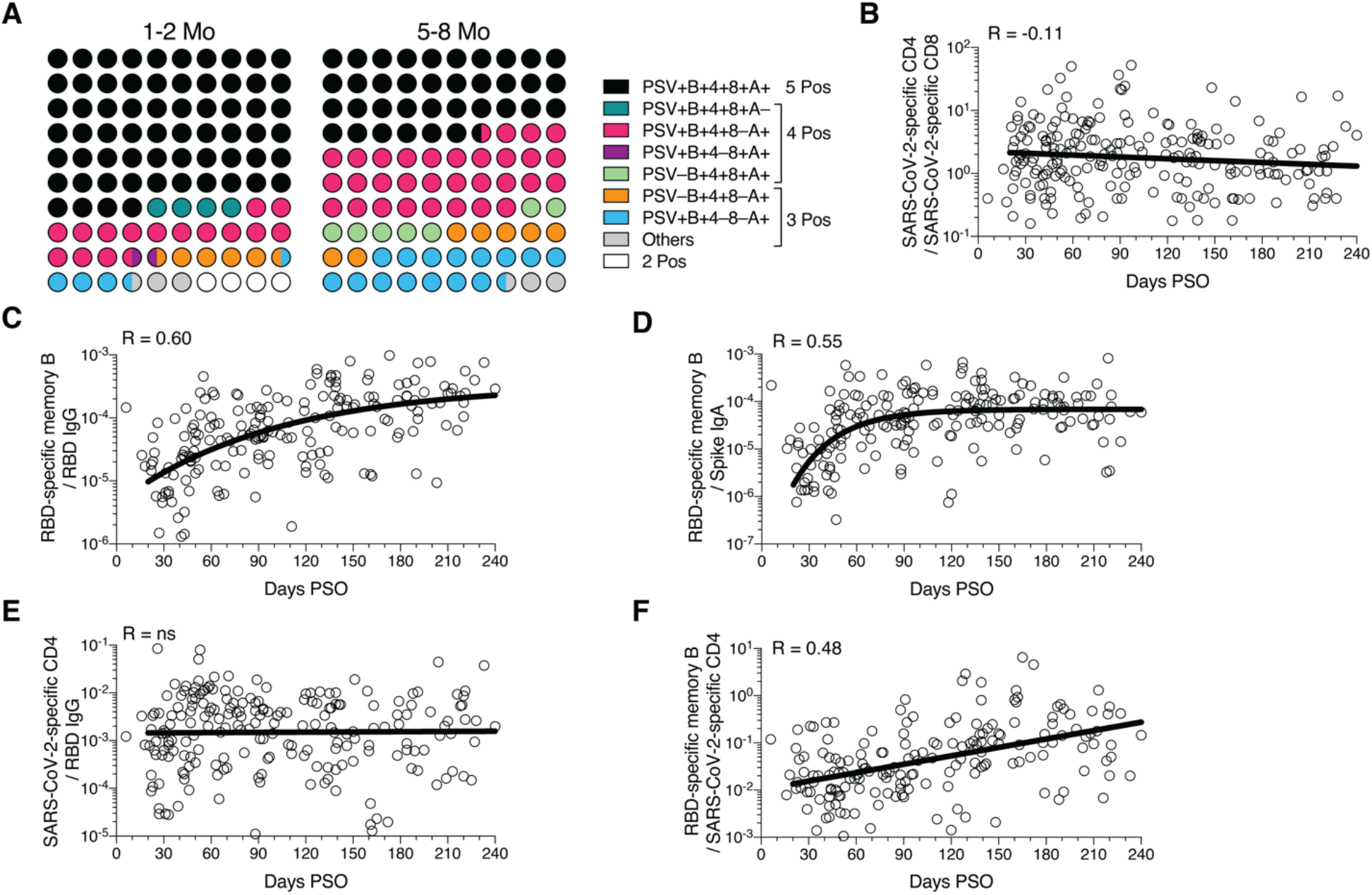
Immune memory relationships. **(A)** Percentage dot plots showing frequencies (normalized to 100%) of subjects with indicated immune memory components during the early (1-2 mo) or late (5-8 mo) phase. “PSV”, PSV-neutralizing antibodies. “B”, RBD-specific memory B cells. “4”, SARS-CoV-2 specific CD4^+^ T cells. “8”, SARS-CoV-2 specific CD8^+^ T cells. “A”, Spike-specific IgA. n = 78 (1-2 mo), n = 44 (5-8 mo). **(B)** The ratio of SARS-CoV-2 specific CD4^+^ T cell frequency relative to SARS-CoV-2 specific CD8^+^ T cell frequency (best-fit simple linear regression line, |R| = 0.11). Three data points are outside the axis limits. **(C)** The ratio of RBD-specific memory B cell frequency (percentage) relative to RBD-specific IgG (pseudo-first order kinetic model, |R| = 0.60). Three data points are outside the axis limits. **(D)** The ratio of RBD-specific memory B cell frequency (percentage) relative to Spike IgA antibodies (pseudo-first order kinetic model, |R| = 0.55). One data point is outside the axis limits. **(E)** The ratio of SARS-CoV-2 specific CD4^+^ T cell frequency relative to RBD IgG antibodies (best-fit simple linear regression line, R = 0.046). Three data points are outside the axis limits. **(F)** The ratio of RBD-specific memory B cell frequency (percentage) relative to total SARS-CoV-2 specific CD4^+^ T cell frequency (best-fit simple linear regression line, |R| = 0.48). One data point is outside the axis limits. For Figure 5H: The ratio of RBD-specific memory B cell frequency (percentage) relative to Spike IgA antibodies (blue curve; best-fit pseudo-first order kinetic curve transformed by ×10^6^), RBD IgG antibodies (orange; best-fit pseudo-first order kinetic curve transformed by × 10^5^) and total SARS-CoV-2 specific CD4^+^ T cell frequency purple; best-fit simple linear regression line transformed by ×10^2^), or the ratio of SARS-CoV-2 specific CD4^+^ T cell frequency relative to SARS-CoV-2 specific CD8^+^ T cell frequency (teal; best-fit simple linear regression line) and RBD IgG antibodies (black; best-fit simple linear regression line transformed by × 10^3^).

**Table S1.** Memory B cell flow cytometry panel.

| Reagents | Source | Identifier | Dilution |
| --- | --- | --- | --- |
| Mouse anti-human CD62L BV615 (clone SK11) | BD Bioscience | Cat# 565219 | 1:200 |
| Mouse anti-human CD19 BUV563 (clone SJ25C1) | BD Bioscience | Cat# 612916 | 1:200 |
| Mouse anti-human FCRL5 (CD307e) BUV615 (clone 509F6) | BD Bioscience | Cat# 751131 | 1:50 |
| Mouse anti-human CD95 BUV737 (clone DX2) | BD Bioscience | Cat# 612790 | 1:200 |
| Mouse anti-human CCR6 BUV805 (clone 11A9) | BD Bioscience | Cat# 749361 | 1:200 |
| Mouse anti-human CD138 BV480 (clone MI15) | BD Bioscience | Cat# 566140 | 1:50 |
| Mouse anti-human IgD BV510 (clone IA6-2) | BioLegend | Cat# 348220 | 1:200 |
| Mouse anti-human IgM BV570 (clone MHM-88) | BioLegend | Cat# 314517 | 1:200 |
| Mouse anti-human CD24 BV605 (clone ML5) | BioLegend | Cat# 311124 | 1:200 |
| Mouse anti-human CD20 BV650 (clone 2H7) | BioLegend | Cat# 302336 | 1:200 |
| Rat anti-human CXCR5 BV750 (clone RF8B2) | BD Bioscience | Cat# 747111 | 1:200 |
| Mouse anti-human CD71 BV786 (clone M-A712) | BD Bioscience | Cat# 563768 | 1:200 |
| Mouse anti-human CD27 BB515 (clone M-T271) | BD Bioscience | Cat# 564642 | 1:200 |
| Mouse anti-human IgA Vio Bright FITC (clone IS11-8E10) | Miltenyi Biotec | Cat# 130-113-480 | 1:400 |
| Mouse anti-human CD3 PerCP (clone SK7) | BioLegend | Cat# 344814 | 1:100 |
| Mouse anti-human CD14 PerCP (clone 63D3) | BioLegend | Cat# 367152 | 1:200 |
| Mouse anti-human CD16 PerCP (clone 3G8) | BioLegend | Cat# 302030 | 1:200 |
| Mouse anti-human CD56 PerCP (clone HCD56) | BioLegend | Cat# 318342 | 1:200 |
| Rat anti-human IgG PerCP/Cyanine5.5 (clone M1310G05) | BioLegend | Cat# 410710 | 1:100 |
| Mouse anti-human CD85j PE/Dazzle 594 | BioLegend | Cat# 333716 | 1:100 |
| Mouse anti-human CD11c PE/Cyanine5 (clone 3.9) | BioLegend | Cat# 301610 | 1:200 |
| Mouse anti-human CD21 Alexa Fluor 700 (clone Bu32) | BioLegend | Cat# 354918 | 1:50 |
| LIVE/DEAD Fixable Blue Stain Kit | ThermoFisher | Cat# L34962 | 1:400 |

**Table S2.** Antibodies utilized in the CD8^+^ and CD4^+^ T cell activation induced markers (AIM) assays

| **Reagents** | **Source** | **Identifier** | **Dilution** |
| --- | --- | --- | --- |
| Mouse anti-human CD45RA BV421 (clone HI100) | BioLegend | Cat# 304130 | 1:50 |
| Mouse anti-human CD14 BUV563 (clone M5E2) | BD Bioscience | Cat# 741360 | 1:100 |
| Mouse anti-human CD19 BUV805 (clone HIB19) | BD Bioscience | Cat# 742007 | 1:100 |
| Fixable Viability Dye eFluor 506 | ThermoFisher | Cat# 65-0866-18 | 1:200 |
| Mouse anti-human CD8a BV650 (clone RPA-T8) | BioLegend | Cat# 301042 | 1:50 |
| Mouse anti-human CD4 BV605 (clone RPA-T4) | BD Bioscience | Cat# 562658 | 1:25 |
| Mouse anti-human CCR7 FITC (clone G043H7) | BioLegend | Cat# 353216 | 1:50 |
| Mouse anti-human CD69 PE (clone FN50) | BD Bioscience | Cat# 555531 | 1:10 |
| Mouse anti-human OX40 PECy7 (clone Ber-ACT35) | BioLegend | Cat# 350012 | 1:50 |
| Mouse anti-human CD137 APC (clone 4B4-1) | BioLegend | Cat# 309810 | 1:25 |
| Mouse anti-human CD3 AF700 (clone UCHT1) | ThermoFisher | Cat# 56-0038-42 | 1:25 |
| Mouse anti-human CD62L BV615 (clone SK11) | BD Bioscience | Cat# 565219 | 1:200 |
| Mouse anti-human CD19 BUV563 (clone SJ25C1) | BD Bioscience | Cat# 612916 | 1:200 |

